# Proline catabolism is key to facilitating *Candida albicans* pathogenicity

**DOI:** 10.1101/2023.01.17.524449

**Authors:** Fitz Gerald S. Silao, Tong Jiang, Biborka Bereczky-Veress, Andreas Kühbacher, Kicki Ryman, Nathalie Uwamohoro, Sabrina Jenull, Filomena Nogueira, Meliza Ward, Thomas Lion, Constantin F. Urban, Steffen Rupp, Karl Kuchler, Changbin Chen, Christiane Peuckert, Per O. Ljungdahl

## Abstract

*Candida albicans*, the primary etiology of human mycoses, is well-adapted to catabolize proline to obtain energy to initiate morphological switching (yeast to hyphal) and for growth. We report that *put1-/-* and *put2-/*- strains, carrying defective <u>P</u>roline <u>UT</u>ilization genes, display remarkable proline sensitivity with *put2*-/- mutants being hypersensitive due to the accumulation of the toxic intermediate P5C, which inhibits mitochondrial respiration. The *put1-/*- and *put2-/-* mutations attenuate virulence in *Drosophila* and murine candidemia models. Using intravital 2-photon microscopy and label-free non-linear imaging, we visualized the initial stages of *C. albicans* cells colonizing a kidney in real-time, directly deep in the tissue of a living mouse, and observed morphological switching of wildtype but not of *put2-/-* cells. Multiple members of the *Candida* species complex, including *C. auris*, are capable of using proline as a sole energy source. Our results indicate that a tailored proline metabolic network tuned to the mammalian host environment is a key feature of opportunistic fungal pathogens.

## Introduction

Proline is the sole proteinogenic secondary amino acid. Its pyrrolidine ring gives it a distinctive role in protein architecture and dynamics. Proline plays an important role in energy generation, stress protection, signaling and redox balance of cells across multiple kingdoms *(1-3)*. Proline is the most abundant amino acid in the extracellular matrix (ECM); collagen, contains ∼23% proline and hydroxyproline combined *(4)*, and mucin, is about ∼13% proline (*5*). In some pathological states, such as cancer and sarcopenia, proline is released in significant quantities as a result of ECM degradation by extracellular proteases (*6, 7*), generating a proline pool that can be assimilated by human cells and the associated microbiome. In eukaryotes, proline catabolism occurs largely in mitochondria, and its complete oxidation can generate ∼30 equivalents of ATP, making proline an excellent energy source (*3, 8, 9*).

*Candida* spp. are the major fungal commensals in humans with *Candida albicans* as the predominant species. *C. albicans* is an opportunistic pathogen capable of causing a spectrum of pathologies ranging from superficial mycoses to life-threatening systemic infections. As a pathogen *C. albicans* must circumvent the host immune response and acquire nutrients to support the bioenergetic demands of infectious growth. Proline is a potent inducer of morphological switching in *C. albicans*, i.e., yeast-to-hyphal growth *(10, 11)*. The inducing properties of proline depends on its catabolism, which stimulates the well-characterized hyphal-inducing Ras1/cAMP/PKA pathway *(10). C. albicans* strains that cannot metabolize proline exhibit defective hyphal growth and reduced survival within macrophages *(10)*. Consistently, strains lacking *GNP2*, encoding the primary proline permease, are unable to filament in the presence of proline and exhibit reduced survival when co-cultured with macrophages *(12)*. Most of the presumed knowledge regarding <u>P</u>roline <u>UT</u>ilization (PUT) in fungi has been extrapolated from studies on the budding yeast *Saccharomyces cerevisiae* (reviewed in *(13)*. The regulatory mechanisms underlying PUT in *C. albicans* have not been well-characterized.

In eukaryotes the catabolic conversion of proline to glutamate is restricted in the mitochondria and is carried out by the concerted actions of proline dehydrogenase (PRODH; EC 1.5.5.2) and Δ^1^-pyrroline-5-carboxylate (P5C) dehydrogenase (P5CDH; EC 1.2.1.88) (reviewed in (*1, 2, 14*). In *C. albicans* PRODH and P5CDH are encoded by *PUT1* (C5_02600W) and *PUT2* (C5_04880C), respectively *(10)*. Put1 uses flavin adenine dinucleotide (FAD) as a co-factor, oxidizing proline to generate P5C and FADH_2_. The electrons from FADH_2_ reduce membrane bound ubiquinone, co-enzyme Q, effectively linking proline oxidation to the mitochondrial electron transport chain (ETC) (*1, 2, 14*). In mammals, PRODH is functionally and physically linked to ETC-Complex II (succinate dehydrogenase) *(14, 15)*. P5C forms a non-enzymatic equilibrium with L-glutamate γ-semialdehyde (GSA), an equilibrium that is pH sensitive; P5C formation is favored with increasing pH *(16)*. Put2 catalyzes the oxidation of GSA to glutamate resulting in the reduction of NAD^+^ to NADH, which is oxidized in an energy conserving manner by NADH dehydrogenase; ETC-Complex I. Glutamate is subsequently converted to α-ketoglutarate (α-KG) by the NAD^+^-dependent glutamate dehydrogenase (GDH; EC 1.4.1.2) releasing ammonia and reduced NADH. In mammalian cells, GDH is localized in the mitochondria and performs an important anaplerotic function by directly feeding the TCA cycle with α-KG *(17)*. In *C. albicans*, however, GDH (Gdh2; C2_07900W) is cytosolic *(18)*. Gdh2 is the key ammonia generating enzyme when *C. albicans* cells utilize amino acids as sole energy source, and it is responsible for the observed alkalization of the extracellular milieu *(18)*.

Previously, we reported that mitochondria-localized proline catabolism induces and energizes hyphal formation in *C. albicans* and that *C. albicans* cells depend on proline catabolism to evade from macrophages *(10)*. Here, we show that multiple pathogenic *Candida* spp. are able to catabolize proline as sole energy source. Using *C. albicans* as a paradigm species, we have carried out a thorough characterization of PUT, focusing on the induction by proline and the bioenergetics of proline-driven virulence. Mitochondrial proline catabolism is tightly regulated to minimize the toxicity of the intermediate P5C, and that PUT is required for virulence of *C. albicans* in *Drosophila* and murine systemic infection models. Finally, using intravital microscopy, we visualized the initial stages of colonization of the kidney *in situ* in a living host and confirm the importance of PUT in the induction of filamentous growth during the early stages of infection.

## Results

### The ability to use proline as sole energy source is common to pathogenic *Candida* species

*C. albicans*, as well as other *Candida* spp., have evolved in the low sugar environment of mammalian hosts and are rarely found living free in nature. *C. albicans* possesses mitochondria with a complete repertoire of electron transport complexes (ETC-Complexes I-V) and is well-adapted to utilize proline as an energy source *(10)*. We examined the possibility that other fungi of the *Candida* pathogenic species complex *(19, 20)* may have evolved the ability to exploit proline as an energy source (Fig. 1). The laboratory *C. albicans* strain SC5314 (dilution 1) and two clinical isolates PLC124 (dilution 2) and MAY7 (dilution 3) *(21)* grew on SP, a minimal synthetic medium containing 10 mM proline as sole energy source (Fig. 1A). In contrast, the haploid S288c (dilution 4) and ∑1278b-derived diploid *(22)* (dilution 5) *S. cerevisiae* strains did not grow on SP, but consistent with their ability to use proline as a nitrogen source they did grow on SPD, which is SP supplemented with 2% glucose (Fig. 1A). *S. cerevisiae* isolated from blood samples obtained from patients with fungal infections were also tested, and similar to the laboratory strains, they were unable to grow on SP.

**Fig. 1.**
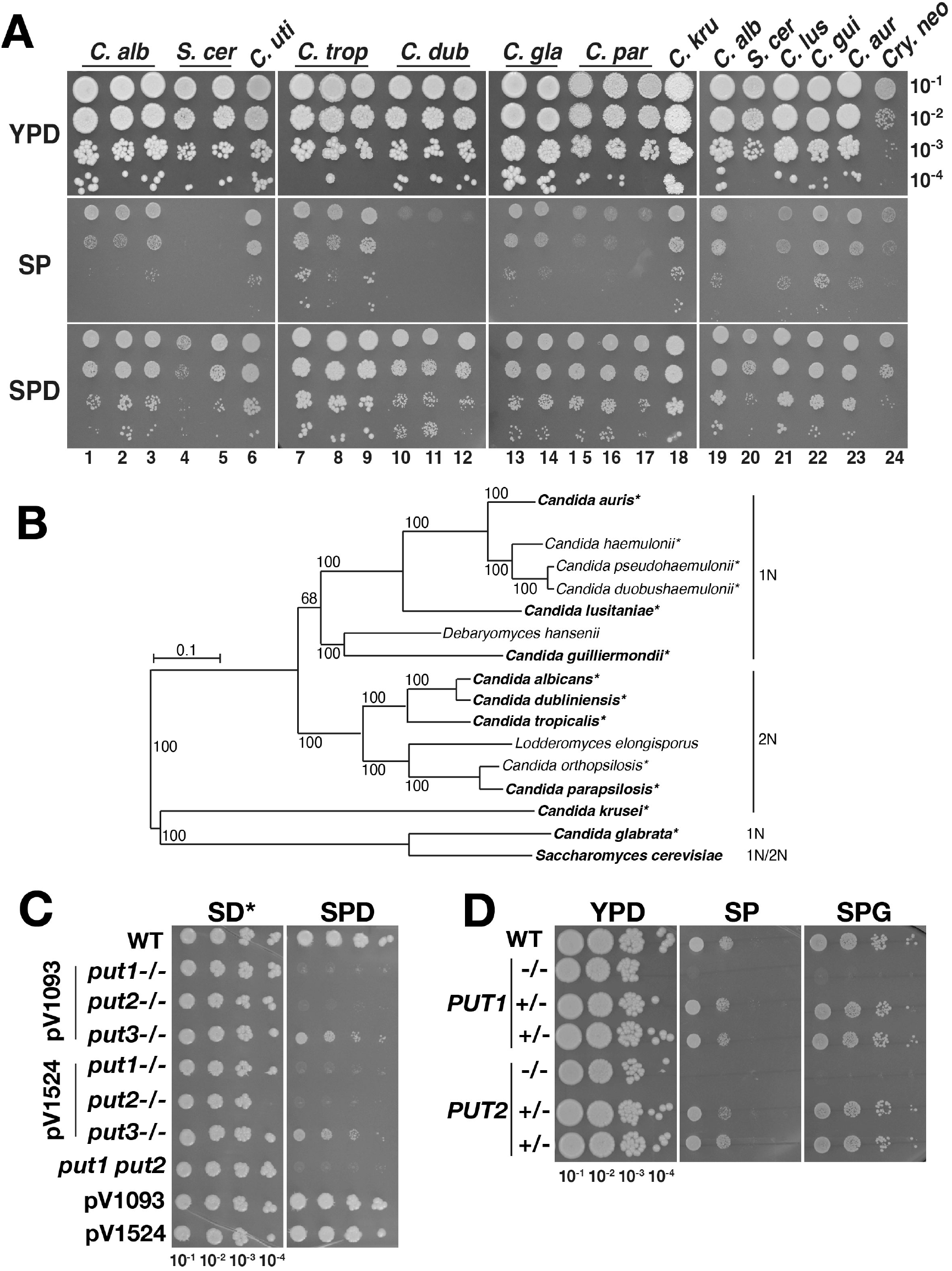
Proline as sole energy source supports growth of diverse pathogenic *Candida* species. (**A**) Proline utilization by members of the *Candida* pathogenic species complex. The ability of the following fungal strains to grow using proline as the sole source of energy was assessed: *C. albicans* (SC5314, PLC124 and MAY7) dilutions 1-3; *S. cerevisiae* (haploid S288c and diploid ∑1278b) dilutions 4 and 5; *C. utilis* (F608) dilution 6; *C. tropicalis*, (SM1541 and ATCC750 (from two different laboratories)) dilutions 7-9; *C. dubliniensis* (SMI718 and Wü284 (from two different laboratories)) dilutions 10-12; *C. glabrata* (CBS138 and Peu927) dilutions 13-14; *C. parapsilosis* (ATCC22019 from three different laboratories) dilutions 15-17; *C. krusei* (ATCC6258) dilution 18; *C. albicans* (SC5314) dilution 19; *S. cerevisiae* (S288c) dilution 20; *C. lusitaniae* (DSM 70102) dilution 21; *C. guilliermondii* (ATCC6260) dilution 22; *C. auris* (B11220) dilution 23; and *Cryptococcus neoformans* (NEQS) dilution 24. All strains were pre-grown on SD from single colonies, washed 2X with H_2_O and resuspended in H_2_O at an OD_600_ ≈ 1. Serial dilutions were prepared and spotted on the indicated medium: YPD, SP (10 mM proline), and SPD (10 mM proline, 2% glucose). The plates were incubated at 30 °C for 48 h and photographed. (**B**) Phylogenetic tree of the *Candida* pathogenic species complex (adapted from (*19*)). The strains in bold text were analyzed for their ability to use proline as sole energy source (**A**); strains with asterisks are known to cause infections in humans. The ploidy, haploid (1N) and diploid (2N), of the strains are as indicated. (**C**) *PUT1* and *PUT2* are essential for proline utilization. Washed cells from YPD pre-cultures were serially diluted and spotted on buffered (pH = 6; 50 mM MES) SD* and SPD, incubated at 30 °C for 4 days. SD* contains a non-standard concentration of ammonium sulfate, 5 mM compared to 38 mM of standard SD. Strains: WT (SC5314); pV1093-derived strains: *put1*-/- (CFG149), *put2*-/- (CFG143) and *put3*-/- (CFG150); pV1524-derived strains: *put1*-/- (CFG154), *put2*-/- (CFG318) and *put3*-/- (CFG156); *put1*-/- *put2*-/- (*put1 put2*, CFG159) derived using both pV1524 and pV1093; and control strains pV1093 (CFG181) and pV1524 (CFG182) carrying the vector without guide RNA. (**D**) Genetic reconstitution to assess the accuracy of CRISPR/Cas9 induced *put1-/-* and *put2-/-* mutations. DNA fragments, amplified from WT (SC5314), that span the CRISPR/Cas9-induced mutations were introduced into strains CFG154 (*put1-/-*) and CFG318 (*put2-/-*). Transformants capable of utilizing proline were selected on SPD media and their growth was assessed on YPD, SP and SPG. Plates were incubated at 30 °C for 4 days. The genotypes of the strains were analyzed by PCR-restriction digest (PCR-RD; Fig. S1B). Reconstituted strains: *PUT1*+/- (CFG379, CFG380), *PUT2*+/- (CFG381, CFG382).

Strikingly, many members of the *Candida* pathogenic species complex were found to utilize proline as sole energy source (Fig. 1A). *C. dubliniensis*, a diploid yeast that is phylogenetically close to *C. albicans* (Fig. 1B), but considerably less virulent *(20)*, exhibited poor growth on SP (dilutions 10-12), whereas *C. tropicalis*, a virulent member of the complex, showed robust growth (dilutions 7-9). Remarkably, *C. glabrata*, a haploid yeast that is phylogenetically close to *S. cerevisiae* and thought to lack an energy-conserving mitochondrial NADH-dehydrogenase (ETC-CI; reviewed in *(23)*) grew on SP (dilutions 13-14). Other members of the complex, such as *C. parapsilosis* (dilutions 15-17), *C. lusitaniae* (dilution 21), *C. krusei* (dilution 18), *C. guilliermondii* (dilution 22), and the more recently characterized multi-drug resistant species *C. auris (19)* (dilution 23) exhibited growth on SP. *C. utilis*, a rare cause of human infection *(24)*, also utilized proline efficiently, consistent with it possessing a functional mitochondrial ETC-CI (dilution 6) *(25)*. The data indicate that the ability to use proline as an energy source is a common attribute among pathogenic *Candida* spp., as well as in *Cryptococcus neoformans* (dilution 24), the latter a pathogenic yeast commonly found in HIV patients *(26)*.

To establish the role of PUT in pathogenic growth we considered *C. albicans* as a paradigm representative of the *Candida* spp. complex. Consistent to our previous report *(10)*, independently generated *put1-/-* and *put2-/-* strains failed to utilize proline as sole nitrogen source (Fig. 1C). Strains transformed with empty vectors (i.e., pV1093 *(27)* and pV1524 *(28)*) retained the ability to grow on SPD. To verify that the Put^-^ phenotypes were linked to modifications at the expected loci, the *put1-/-* and *put2-/-* strains were transformed with wildtype *PUT1* or *PUT2* fragments and Put^+^ revertants were selected on SPD (Fig. S1A). The heterozygous revertants fully regained the ability to grow using proline as sole carbon and nitrogen source (SP and SPG; Fig. 1D); restoration of the wildtype alleles was confirmed by PCR (Fig. S1B). We also inactivated *PUT1* and *PUT2* in a *cph1*Δ/Δ *efg1*Δ/Δ non-filamenting *C. albicans* strain *(29)* by CRISPR and in *C. glabrata* using a modified SAT1-flipper technique *(30)*, which completely abrogated their ability to grow on SPD (Fig. S1C).

### Proline catabolism is induced by proline in a Put3-dependent and -independent manner

In *C. albicans*, as in *S. cerevisiae*, proline catabolism is induced by the presence of proline and due to being mitochondrial localized is sensitive to glucose repression (reviewed in (*8*)) (Fig. 2A). In contrast to *S. cerevisiae, PUT1* and *PUT2* expression in *C. albicans* is not under nitrogen catabolite repression; their expression is induced by proline in a Put3-dependent manner even in cells grown in the presence of 38 mM ammonium *(10, 31)*. Proline is transported into mitochondria by an undefined process and the glutamate formed is directly or indirectly transported to the cytoplasm where it is deaminated to α-ketoglutarate in a reaction catalyzed by cytoplasmic glutamate dehydrogenase (Gdh2) *(18)*. To better assess the mechanisms regulating proline catabolism, we created a reporter strain co-expressing Put1-RFP, Put2-HA, and Gdh2-GFP (Fig. S1D), which enabled the simultaneous analysis of the expression of the PUT enzymes and Gdh2 by immunoblot. Consistent with our previous work *(18)*, subcellular fractionation and microscopy confirmed that the tagged constructs localized correctly; Put1-RFP and Put2-HA are clear mitochondrial components and Gdh2-GFP localizes to the cytoplasm (Fig. 2B).

**Fig. 2.**
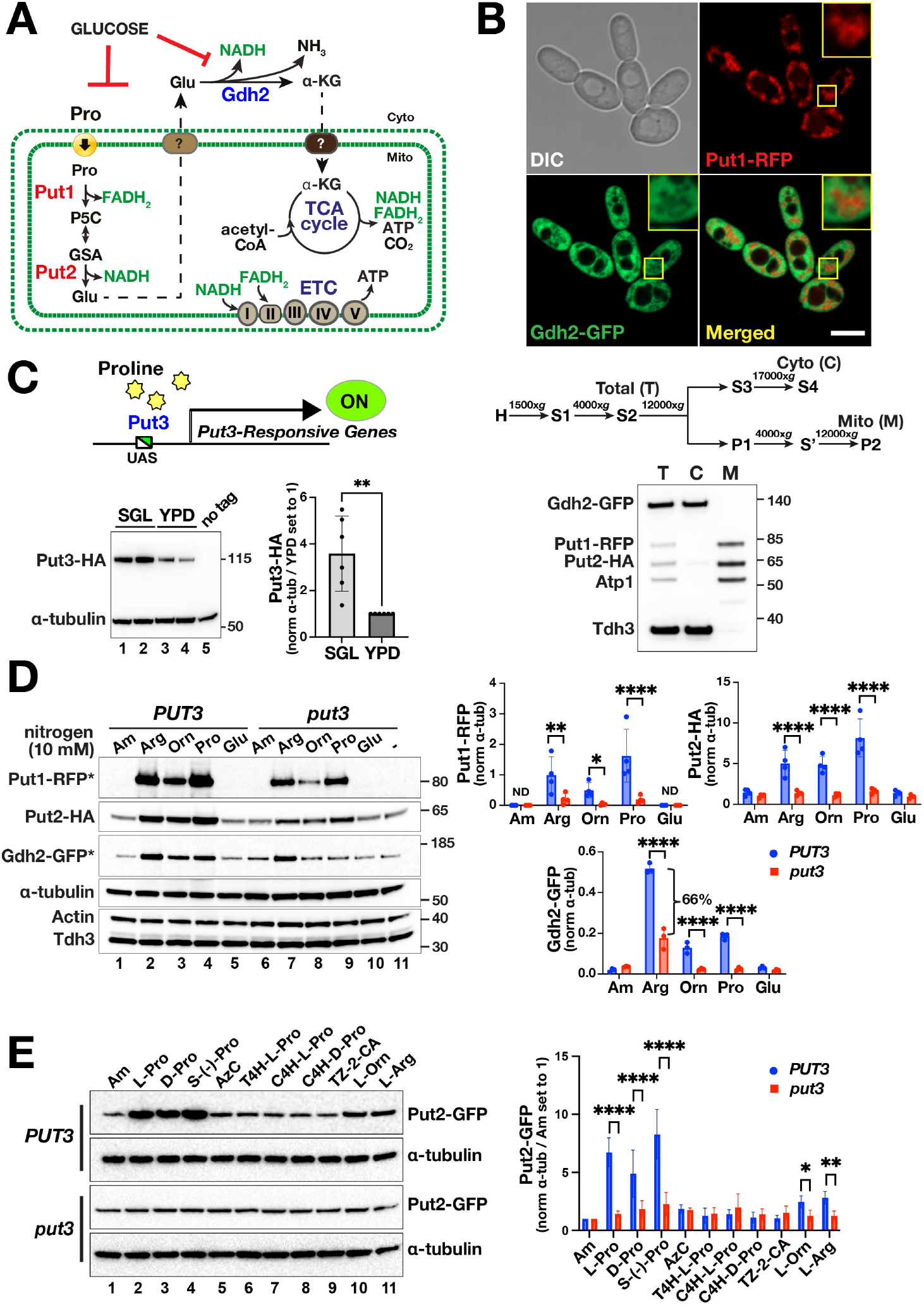
Proline catabolism is induced by proline in a Put3-dependent and -independent manner. (**A**) Scheme of proline catabolism in *C. albicans*. Proline is transported into mitochondria by an unknown process and converted to glutamate in two enzymatic steps, catalyzed by Put1 and Put2, generating reduced electron carriers that are oxidized by the electron transport chain (ETC) resulting in the generation of ATP. Glutamate is deaminated in the cytoplasm by Gdh2. The presence of glucose represses mitochondrial functions and leads to reduced Gdh2 expression. (**B**) Microscopy and subcellular fractionation of proline catabolic enzymes. Upper panels; representative confocal (Airyscan) image of strain CFG407 grown in YPG for 4 h. Lower panels; homogenized cell extracts prepared from CFG433, pre-grown in YPD and shifted to YPG 4h, 37° C, were analyzed according to the indicated centrifugation scheme; homogenized cell extracts (H), supernatant (S) and pellet fractions (P). The Total (T) (S2), cytosolic (C) (S4) and mitochondrial (M) (P2) fractions were analyzed by immunoblotting. Put1-RFP (∼ 81.9 kDa) and Put2-HA (∼ 69.8 kDa) co-fractionate with the mitochondrial marker (Atp1, ∼ 59 kDa). Whereas Gdh2-GFP (∼ 141 kDa) co-fractionates with the cytosolic enzyme, Tdh3 (∼ 35.8 kDa). The micrographs and subcellular fractionation are consistent and demonstrate the clear spatial exclusivity of Put1-RFP and Gdh2-GFP signals. (**C**) Put3 expression is influenced by the growth medium. Upper panel, schematic diagram of proline-dependent activation of Put3-responsive genes. Lower panel (left), strains expressing Put3-HA (CFG187, lanes 1 and 3; CFG188, lanes 2 and 4) were grown to log phase in SGL and YPD as indicated and processed for immunoblotting. Lower panel (right), the relative strength of the immunoreactive Put3-HA signals were determined; the signals were normalized to α-tubulin and to the levels in YPD set to 1 (Ave.±SD, n = 6; \*\**p* <0.01 by student *t*-test). (**D**) Put3-dependent and -independent induction of PUT enzymes. Left panels, immunoblot analysis of extracts prepared from exponentially growing cultures of *PUT3*+/+ (*PUT3*, CFG441) and *put3-/-* (*put3*, CFG443) in SGL 1 h after the addition of 10 mM of the indicated nitrogen sources. The Put1-RFP and Gdh2-GFP signals (*) were enhanced for display via the high slider in Image Lab (BioRad). Right panels, the relative strengths of the immunoreactive bands were determined; the signals were normalized to α-tubulin (Ave.±SD, n ≥ 3; \*\*\*\**p* <0.0001, \*\**p* <0.01, \**p* <0.05 by 2-way ANOVA with Sidak’s post hoc test). Note that Put1-RFP is not detected (ND) in the absence of proline. (**E**) Put3 exhibits specificity for proline. Left panels, immunoblot analysis of extracts prepared from exponentially growing cultures of *PUT3+/+* (*PUT3*, CFG259) and *put3-/-* (*put3*, CFG301) in SGL 1 h after the addition of 10 mM of the indicated compounds: Ammonium sulfate (Am); L-proline; D-proline; S-(-)-proline; Azetidine carboxylate (AzC); T-4-hydroxy-L-proline; C-4-hydroxy-L-proline, C-4-hydroxy-D-proline; Thiazolidine-2-carboxylic acid; L-ornithine; and L-arginine. The extracts were processed as in (D). The relative strength of the immunoreactive Put2-GFP signals were determined; the signals were normalized to α-tubulin and to the levels in Am set to 1 (Ave.±SD, n = 4) and were analyzed by 2-way ANOVA with Sidak’s post hoc test (\*\*\*\**p* <0.0001, \*\**p* <0.01, \**p* <0.05).

Put3 is a large transcription factor in *S. cerevisiae* that contains an N-terminal Zn(II)_2_Cys_6_ binuclear cluster DNA binding domain and a C-terminal domain that undergoes a proline-dependent conformational change that activates *PUT* gene expression *(32, 33)*. Although in *C. albicans*, the highly conserved Put3 (*PUT3*; C1_07020C) homologue facilitates proline-responsive transcriptional activation of *PUT1* and *PUT2 (31)*, inactivation of *PUT3* did not fully abolish the growth of *C. albicans* on SPD (Fig. 1C), indicating Put3-independent mechanisms operate. Strikingly, Put3-HA expression is not repressed by ammonium but is sensitive to glucose; Put3-HA is readily detected in cells grown in SGL containing 38 mM ammonium and glycerol and lactate as carbon sources, whereas expression is significantly lower in YPD containing 2% glucose (Fig. 2C). This aligns with *PUT2* expression being lower in cells grown in high glucose media *(10)*.

We assessed the expression of Put1-RFP, Put2-HA, and Gdh2-GFP in *PUT3*+/+ (CFG441) and *put3-/-* (CFG443) cells 1 h after different nitrogen sources (10 mM) were added to exponentially growing cultures in SGL (Fig. 2D). Put1-RFP was not detected in control cultures in the absence of additional nitrogen (-; lane 11) or with ammonium (Am; lane 1). In contrast and as reported *(10)*, basal Put2-HA levels were expressed in both strains (lanes 1,11). All three reporters in *PUT3*+/+ cells were significantly induced upon the addition of 10 mM proline (lane 4). Arginine and ornithine, which can be metabolically converted to proline *(10)*, also induced their expression (lanes 2,3). The lower induction of Put1-RFP in ornithine spiked cultures is likely due to the SPS-sensor dependency of its uptake (lane 3) *(10)*. Previous work, exploiting ChIP-Seq to identify Put3-regulated genes in *C. albicans*, did not identify *GDH2* as a target; *(31)* however, these studies were conducted using cells grown in the presence of high glucose (YPD), a condition that represses *PUT3* expression (Fig. 2C) and *GDH2* expression (*18*). The proline-dependent Put1 expression occurred independent of Put2 (Fig. S2C). Unexpectedly, the addition of glutamate, which in mammals is a precursor of proline biosynthesis *(2, 14)*, did not induce reporter expression (Fig. 2D; Fig. S2A, C) nor did glutamine (Fig. 2SA). Notedly, proline induced the expression of Put1-RFP and Put2-HA in *put3*-/- cells, albeit to a significantly lower level than in the *PUT3+/+* strain (Fig 2D, lanes 4,9; Fig. S2B). The low, but significant Put3-independent expression is consistent with the ability of *put3-/-* strains to grow on SPD, when proline merely serves as a nitrogen source (Fig. 1C). Interestingly, arginine, but not proline, partially induced Gdh2-GFP in *put3*-/-, albeit at levels ∼ 66% lower than that observed in *PUT3+/+* (Fig. 2D, compare lanes 2 with 7). Apparently, additional proline- and arginine-sensitive factors contribute to the expression of the reporter constructs.

We tested the substrate specificity of Put3 by examining the capacity of proline and different proline analogs to induce Put2-GFP in *PUT3+/+* (*PUT3*, CFG259) and *put3-/-* (*put3*, CFG301) strains (Fig. 2E). To avoid potential indirect effects of analog toxicity, we monitored expression levels 1 h after addition of the compounds. The ability of Put3 to induce Put2 exhibited high specificity for L-proline, S-(-)-proline (a racemic pure form of L-proline) and D-proline (lanes 2-4), whereas none of the analogs (lanes 5-9), even the closely-related hydroxyproline compounds (lanes 6-8), induced expression. As previously observed (Fig. 2D) and consistent with being catabolized to proline, the addition of ornithine and arginine resulted in the induction of Put2-GFP (lanes 10-11). Similar results were obtained using the triple reporter *PUT3+/+* (CFG441) and *put3-/-* (CFG443) strains (Fig. S2B).

### Proline is toxic in cells unable to catabolize it

We considered the possibility that mutations inactivating proline catabolism could affect mitochondrial function and assessed growth under repressing (glucose) and non-repressing (glycerol, lactate) conditions. The mutants *put1-/-, put2-/-* and *gdh2-/-* grew at indistinguishable rates independent of the absence or presence of 10 mM proline under repressing conditions with2% glucose (SD) (Fig. 3A, upper panel). However, when grown under non-repressing conditions (SGL), the addition of 10 mM proline resulted in striking growth inhibition of *put1-/-* and *put2-/-* cells (Fig. 3A, lower panel). In the absence of proline, *put1-/-* and *put2-/-* mutants grew as well as wildtype, indicating that proline inhibits growth only when cells cannot catabolize it. Cells lacking Put2 exhibited extreme hypersensitivity to proline, only a minimal increase in OD_600_ was observed after 20 h. In contrast, consistent with proline being an excellent energy source, the addition of proline enhanced the growth of the wildtype and *gdh2-/-* strains. Similar results were obtained on solid media (Fig. S3A) and in and on amino acid-rich complex media (Fig. S4A, B). Notedly, on YPD containing 2% glucose, the *put* mutants grew similarly to wildtype. However, when grown in or on YPG and YPL containing non-fermentable carbon-sources, *put2-/-* mutant strains exhibited poor growth. The *put2-/-* cells grown in YPG exhibited defects in cell separation forming trimera (Fig. S4A, see inset), a phenotype associated with stress (*34*). The growth-inhibitory effects of proline are dependent on non-fermentative conditions when mitochondrial respiration is required. The lack of proline inhibition of cells grown in high glucose media is due to glucose repression of mitochondrial activity *(10)* (reviewed in *(8)*). Consistent with this notion, Put1 and Put2 protein levels are elevated in wildtype cells 1 h after the addition of 10 mM proline in SGL and to a lesser extent in SD (Fig. S3B), but in SD, due to repression of mitochondrial function, proline does not inhibit growth. The proline hypersensitivity exhibited by *put2-/-* mutants is partially rescued by the introduction of *put1-/-* but not *gdh2-/-* (Fig. 3A); consequently, the primary growth inhibitory effect of proline is linked to catabolic intermediates formed by Put1 and that are metabolized further by Put2, i.e., either P5C or GSA. We further tested the inhibitory effects of proline in strains grown on synthetic medium containing a non-repressing level of glucose (0.2%) and glutamate as nitrogen source (Fig. S3D), and consistently, we observed the same epistatic relationship between *put1-/-* and *put2-/-*.

**Fig. 3.**
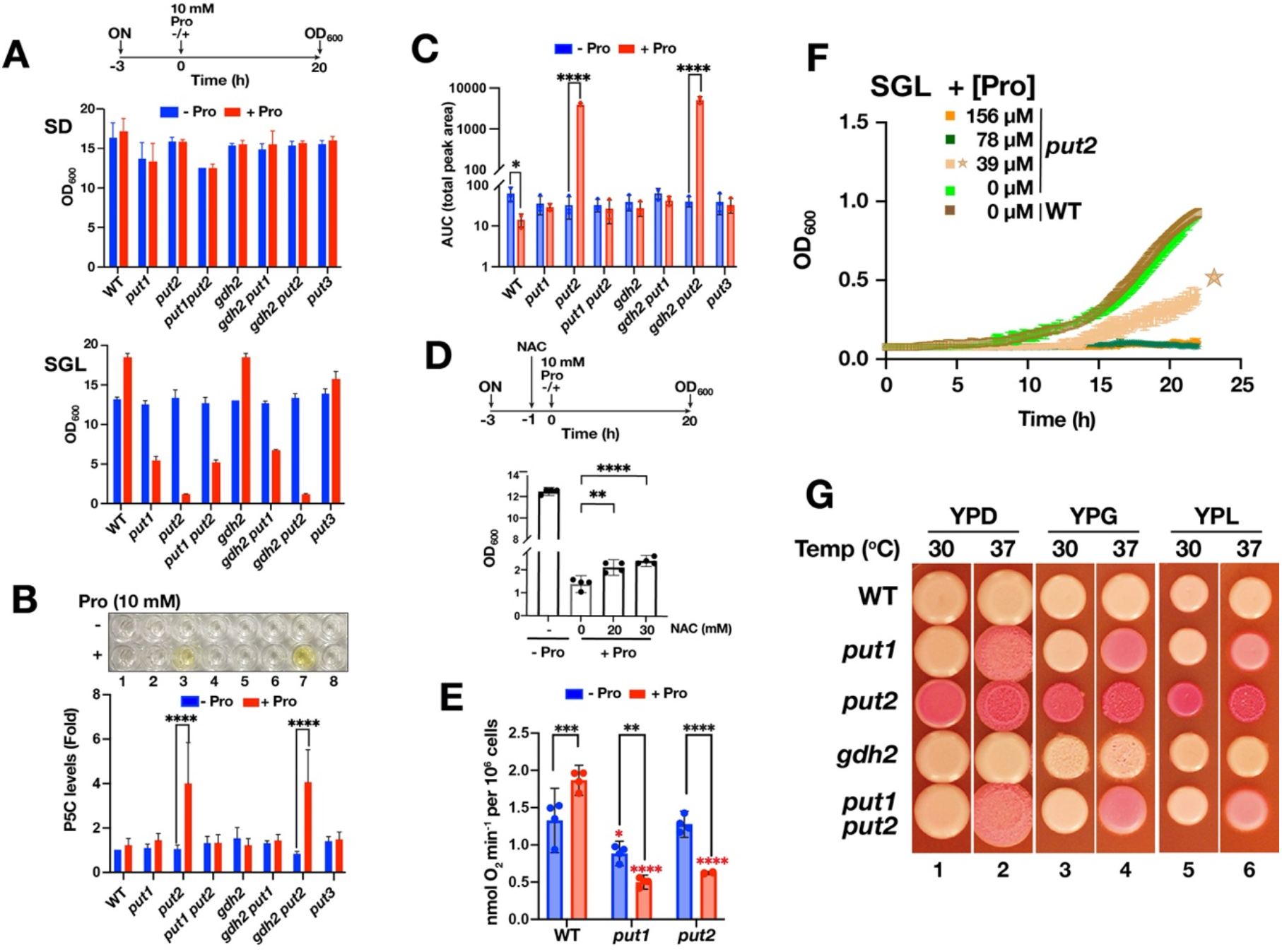
Proline is toxic in cells unable to catabolize it. (**A**) Overnight (ON) cultures of strains with the indicated genotypes in SD and SGL were used to inoculate fresh media and incubated 3 h shaking to obtain exponentially growing cultures. At t=0 (OD_600_ = 1) the cultures were split into two tubes and one tube received 10 mM proline (red) and the other an equal volume of H_2_O (blue). Growth was measured (OD_600_) after incubation with shaking for 20 h at 30 °C. Strains used: WT (SC5314); *put1* (CFG154), *put2* (CFG318); *put3* (CFG156); *put1 put2* (CFG159); *gdh2* (CFG279); *gdh2 put1* (CFG364); *gdh2 put2* (CFG366). Null strains are indicated in lower case letters (e.g., *put1* = *put1-/-*). Results presented (Ave.±SD; n=3) were analyzed by 2-way ANOVA with Sidak’s post hoc test). All pairwise comparisons for SGL have *p*<0.0001 except *put3* (*p*<0.001). (**B**) Inactivation of *PUT2* leads to the accumulation of P5C in the presence of proline. Ten mM proline or an equal volume of H_2_O was added to exponentially growing cultures of the strains as in (A) in SGL and the levels of P5C were determined after 2 h. The data are presented as fold change of P5C levels relative to wildtype strain grown without proline (set to 1). Results presented (Ave.±SD; n=5) were analyzed by 2-way ANOVA with Sidak’s post hoc test (**** *p*<0.0001). (**C**) Inactivation of *PUT2* generates elevated ROS in the presence of proline. Strains as in (A) were grown as in (B) and the levels of ROS was determined 6 h after the addition of 10 mM proline or an equal volume of H_2_O. The data are presented as area under the curve (AUC) (Ave.±SD; n=3) and analyzed by 2-way ANOVA with Sidak’s post hoc test (**** *p*<0.0001, \**p* <0.05). (**D**) Proline hypersensitivity of *put2* mutants is largely independent of ROS accumulation. Strain CFG318 grown as in (A) in SGL, and 1 h prior to proline addition cultures were diluted 1:1 with SGL containing freshly dissolved n-acetylcysteine (NAC) to the indicated concentration. Growth was measured (OD_600_) after incubation with shaking for 20 h at 30 °C. Results presented (Ave. with 95% CI) were analyzed by one-way ANOVA (*p*=0.0002) with Dunnett’s post hoc test relative to no NAC (\*\*\*\**p*<0.0001, \*\**p* <0.01). (**E**) Proline inhibits respiration in *put* mutants. Proline (10 mM; + Pro) or an equal volume of H_2_O (-Pro) was added to exponentially growing cultures of *put1* (CFG154) and *put2* (CFG318) in SGL 4 h prior to measuring oxygen consumption. Data are presented as mean with 95% CI (n = 4) and were analyzed either by 2-way ANOVA (effect of proline addition; black asterisks) or one-way ANOVA (effect of mutation on respiration; red asterisks) with multiple comparison test (\*\*\*\**p*<0.0001, \*\*\**p* <0.001, \*\**p* <0.01, \**p* <0.05). (**F**) Cells carrying *put2* mutations are hypersensitive to submillimolar concentrations of proline. The growth of *put2-/-* (CFG318) was monitored in a TECAN microplate reader in a 100 μl of SGL culture ± indicated submillimolar concentrations of proline. The starting OD_600_ ≥ 0.05. The data as in **Fig. S3F**, is presented as Ave.±SD (n=4). (**G**) Cells lacking the capacity to catabolize proline exhibit enhanced death. Exponential cultures of WT (SC5314), *put1* (CFG154), *put2* (CFG318), *gdh2* (CFG279), and *put1 put2* (CFG159) in YPD were harvested, washed and resuspended in H_2_O at an OD_600_ ≈ 1, and 5 μl were spotted on the indicated medium containing the viability indicator Phloxine B (10 μg/ml). The plates were incubated for 3 days at the indicated temperature and photographed. Phloxine B accumulates in dead cells.

### P5C mediates a respiratory block

P5C is an unstable intermediate postulated to inhibit mitochondrial respiration and concomitantly to enhance ROS formation *(35)*. To test this notion, we used an improved protocol relying on complex formation between P5C and o-aminobenzaldehyde (o-AB) *(36)* to quantify P5C in *C. albicans* strains grown in SGL with and without proline. Initially, we had difficulty reproducibly measuring P5C; success was achieved by direct lysis of whole cells under acidic conditions (TCA) to stabilize P5C. The results confirmed that in the presence of proline, P5C levels are significantly elevated in *put2-/-* cells but not in *put1-/- put2-/-* double mutants; the yellow P5C/o-AB complex became readily observable (Fig. 3B). Next, we measured ROS production using the luminol-HRP system. In the absence of proline, ROS levels were similar. However, upon proline addition, the ROS production increased dramatically in *put2-/-* and *put2-/- gdh2-/-* strains, but not in the *put1-/- put2-/-* double mutant (Fig. 3C). These results are consistent with Put1 acting upstream of Put2. To test if the increased ROS levels account for the proline hypersensitivity of *put2-/-*, we grew the cells in the presence of well-characterized ROS scavengers, e.g., n-acetylcysteine (NAC). Although we could see a slight dose-dependent increase in growth in the presence of NAC relative to the control (no NAC), NAC failed to rescue the growth to a level comparable to cells grown in the absence of proline (Fig. 3D). Likewise, cell permeable ROS scavengers like Mito-TEMPO *(37)* or TIRON *(38)* did not restore growth (Fig. S3E). We conclude that the proline hypersensitive phenotype of *put2-/-* is linked to a P5C-mediated respiratory block and not due to secondary effects resulting from ROS accumulation.

To test this notion directly, we measured oxygen consumption in *put1-/-* and *put2-/-* cells grown in the presence or absence proline. Relative to wildtype, the respiration rate of *put2-/-* was reduced significantly when proline was present (Fig. 3E). Using purified P5C, Nishimura et al. showed that *S. cerevisiae* is sensitive to P5C with an IC_50_ value of 23.8 μM (*35*). We determined the lowest level of proline capable of inhibiting growth of *put2-/-* cells and found that 39 μM proline significantly inhibited growth (Fig. 3F and Fig. S3F), which is 7-fold lower than the mean physiological level of proline in human plasma (276 μM) *(39)*. Interestingly, although Put1 expression is low in the absence of proline (Fig. 2D), we observed a significant reduction in oxygen consumption in *put1-/-* cells even in the absence of proline, and the rate of oxygen consumption was reduced even further when proline was added (Fig. 3E). This observation suggests that proline itself is growth inhibitory, and the capacity to metabolize proline is requisite to alleviate this effect.

### Defective proline utilization is linked to cell death

Next, we assessed whether the growth inhibitory effects arising from incomplete proline catabolism had additional and perhaps lethal consequences. We assayed viability using Phloxine B (PXB), which has been used to qualitatively assess cell death in yeast colonies; PXB crosses biological membranes and accumulates in cells that lack the metabolic energy to extrude it *(40)*. Consequently, colonies with dead cells accumulate PXB and take on a red appearance, the degree of redness reflects the number of dead cells. As shown in Fig. 3G, colonies derived from the *put2-/-* mutant cells accumulated significant amounts of PXB on complex YP media containing glucose (YPD) and with non-fermentable carbon-sources glycerol (YPG) and lactate (YPL). Macro colonies derived from *put1-/-* and *put1-/- put2-/-* strains clearly accumulate PXB when grown at 37 °C, whereas PXB is restricted to the colony periphery when grown at 30 °C. PXB did not accumulate in *gdh2-/-* mutants on any of the media and temperatures tested (Fig. 3G). We observed the same levels of PXB accumulation in Put^-^ deficient stains constructed in the non-filamenting *cph1*Δ/Δ *efg1*Δ/Δ background (*29*) (Fig. S4C). We attempted to obtain a quantitative assessment of cell death using propidium iodide (PI) staining, however, we found that *put* mutants flocculate in liquid culture, precluding measurements by FACS. The degree of flocculation increases as cultures become saturated (Fig. S4D). By microscopy we observed an increased number of PI^+^ cells in 48 h-old YPD cultures of *put1-/-* and *put2-/-* but not *gdh2-/-* strains (Fig. S4F), which correlates with the relief of glucose-mediated repression of mitochondrial functions. Consistent with the pronounced growth inhibition by proline under non-repressing respiratory conditions (SGL) (Fig. 3A), we interpret that the increased cell death of *put2-/-* and *put1-/-* mutants on YPG and YPL media as being the consequence of derepressed mitochondrial activity. Interestingly, the colonies derived from cells carrying *put2-/-* become distinctly yellow in appearance upon prolonged incubation on YPD, presumably due to P5C accumulation (Fig. S4G).

### Proline catabolism is required for invasive growth

We recently showed that *C. albicans* cells rely on proline catabolism to induce and energize hyphal growth in phagosomes of engulfing macrophages *(10)*. Relative to the wildtype, *put* mutants exhibit defects in producing long invasive hyphal filaments around the periphery of macro colonies growing on Spider medium (Fig. 4A), which contains mannitol as primary carbon source *(41)*. The diminished filamentous growth was more pronounced in *put2-/-* (Fig. 4A). We developed a collagen invasion assay to further test the role of proline catabolism in powering invasive growth (Fig. 4B). Cells, applied on top of a collagen plug (Purecol EZ in DMEM/F12 medium) in a transwell, were monitored for their ability to induce filamentous growth, pass through the membrane at the bottom of the transwell and to reach the recovery medium (DMEM). The results were clear, the invasion process was dependent on the ability of cells to catabolize proline. In contrast to wildtype cells, the *put* mutants where not detected in the recovery medium. As control, a non-filamenting *cph1*Δ/Δ *efg1*Δ/Δ strain was used, and as expected, did not invade the collagen matrix. Next, we performed pairwise competition experiments mixing equal numbers of wildtype and mutant cells before applying them on the collagen plug (Fig. 4C). At the end of the 14-day incubation period we quantified the total number cells in the plug and recovery medium, and determined the number of wildtype (WT) cells based on their ability to grow on SPD. For competition with *cph1*Δ/Δ *efg1*Δ/Δ control cells, we quantified wrinkled (WT phenotype) and smooth colonies (mutant phenotype) on Spider agar incubated at 37 °C for 3-4 days. Starting with an input ratio of ∼ 1:1 (WT:*put*); all cells in the recovery medium were wildtype and in the collagen plug the *put* mutants were overgrown by wildtype (Fig. 4C). By contrast, *cph1*Δ/Δ *efg1*Δ/Δ cells, which are unable to form hyphae but capable of using proline, were recovered in the collagen plug almost at the same proportion as wildtype. The results indicate that proline catabolism is required by *C. albicans* to grow and invade collagen matrices.

**Fig. 4.**
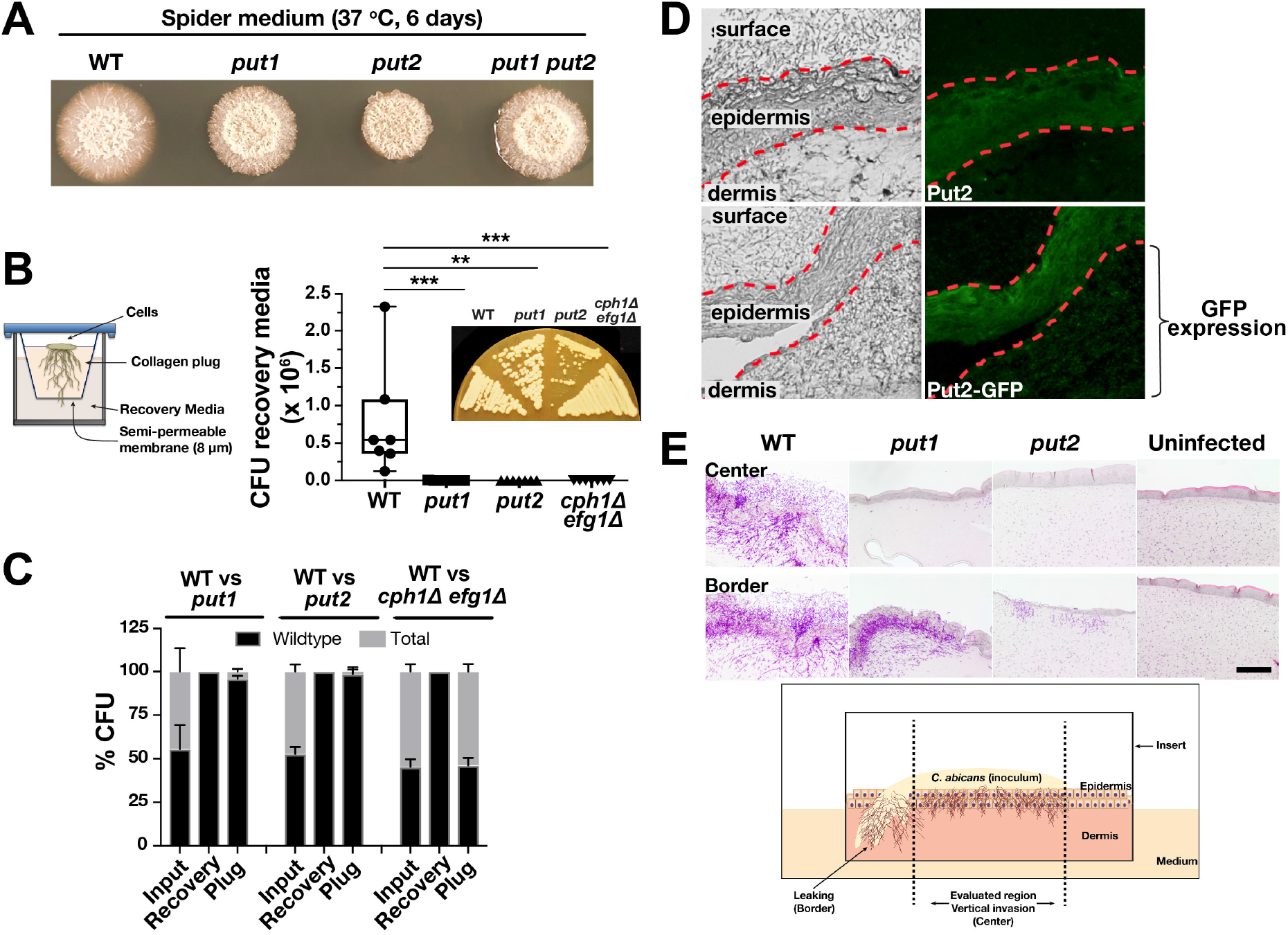
Proline catabolism is required for invasive growth. (**A**) Cells from overnight cultures of WT (SC5314), *put1* (CFG154), *put2* (CFG318), and *put1 put2* (CFG159) in YPD were harvested, washed, resuspended in PBS (OD_600_ ≈ 1) and 5 μl were spotted on Spider medium, incubated 6 days at 37 °C, and photographed. Null strains are indicated in lower case letters (e.g., *put1* = *put1-/-*). (**B**) Schematic diagram of the collagen invasion assay. WT (SC5314), *put1* (CFG154), *put2* (CFG318), and *cph1*Δ/Δ *efg1*Δ/Δ (CASJ041) strains added on the top of the collagen gel (PureCol EZ) in a transwell insert, incubated 14 days at 37 °C after which the CFU in the recovery media was determined (see Methods). Box and whiskers plot derived from seven biological replicates analyzed by Kruskal-Wallis test (\*\*\*\**p*<0.0001) followed by Dunnett’s post hoc test (\*\*\**p* <0.001, \*\**p*<0.01). (Inset) Cells atop the collagen matrix after the 14-day growth were restreaked on YPD and then incubated for 2 days at 30° C. (**C**) The *put* mutants display reduced fitness compared to wildtype in a competitive collagen invasion assay. Cells as in (**B**) were mixed at a 50:50 ratio as indicated. The mixture of WT:mutant cells were added on top of the collagen gel and incubated as in (**B**). Cells were recovered from input, recovery media and plug (see Methods). The genotype of the cells in the input, recovery media and plug were inferred by growth based-assays on SPD or Spider media to determine the WT:mutant ratio. (**D**) Proline catabolism is activated during invasive growth into reconstituted human skin. *PUT2* (PLC016) and *PUT2-GFP* (CFG219) cells were applied on the top of the stratified epidermal layer as indicated, and 2 days post infection, invasive growth was monitored by fluorescence microscopy. The dashed red lines demarcate the surface and dermal faces of the epidermal layer. Keratinocytes in the epidermal layer exhibit autofluorescence. Note the enhanced Put2-GFP signal in the epidermal and underlying dermal layers. (**E**) Proline catabolism is required for invasive growth through reconstituted human skin. Upper panels, Periodic Acid-Schiff (PAS) staining of skin model 2 days after infecting with WT (SC5314), *put1* (CFG154) and *put2* (CFG318) as indicated. Lower panel, schematic diagram of the infection model depicting the center and border areas. Compared to WT, *put1*and *put2* cells exhibit essentially a non-invasive phenotype in the center, similar to the uninfected control, and the greatly reduced capacity of *put2* cells to grow invasively is clearly evident.

To test these findings in a more complex and physiologically relevant assay system, we used an *in vitro* 3D skin model supplemented with functional immune cells as an infection platform *(42)*. First, we assessed whether proline is actively utilized by wildtype cells during the invasion process. To address this, we used a wildtype strain expressing Put2-GFP as a reporter strain (Fig. 4D). Compared to the untagged Put2 strain that served to control for background fluorescence, significant GFP-dependent signal was observed in fungal cells invading the dermis layer. Consistent with the collagen invasion assay, the *put* mutants exhibited lower invasiveness compared to wildtype (Fig. 4E) and filamenting fungal cells were not observed in the center of the skin model. However, *put1-/-* but not *put2-/-* cells filamented at the edge of the skin model, likely due to direct exposure to the filament inducing cell culture medium (see schematic of the model; Fig. 4E). These results highlight the critical role of proline catabolism and the importance of Put2 to induce invasive growth.

We directly compared the ability of physiologically relevant sources of proline, other than collagen, e.g., serum albumin, mucin (from porcine stomach), and hemoglobin, to induce the expression of Put1 and Put2 (Fig. S5). Contrary to our expectation, collagen did not result in elevated expression of Put1, suggesting that free proline is not immediately accessible to *C. albicans*. Interestingly, mucin, a major glycoprotein that lines mucosal membranes, e.g., the gut where *C. albicans* systemic infections can originate *(43)*, robustly induced Put1 and Put2 expression.

### *C. albicans* virulence is related to the ability to catabolize proline

The virulence of *put* strains was assessed in *Drosophila melanogaster* and mouse infection models. We first used an improved mini-host model exploiting *Bom*^*Δ55C*^ flies *(44)*, which lack 10 Bomanin genes on chromosome 2 encoding secreted peptides with antimicrobial property *(45)*. As shown in the survival curves, the *put* mutants displayed greatly diminished virulence compared to wildtype (Fig. 5A). Six days following infection, flies infected with the *put1-/-, put3-/-* or *put1-/- put2-/-* showed general survival rates of 70%, 50%, and 65%, respectively. In comparison, only 15% of the flies infected with wildtype survived. Strikingly, the *put2-/-* mutant was avirulent, survival exhibited precise overlap with the PBS control.

**5.**
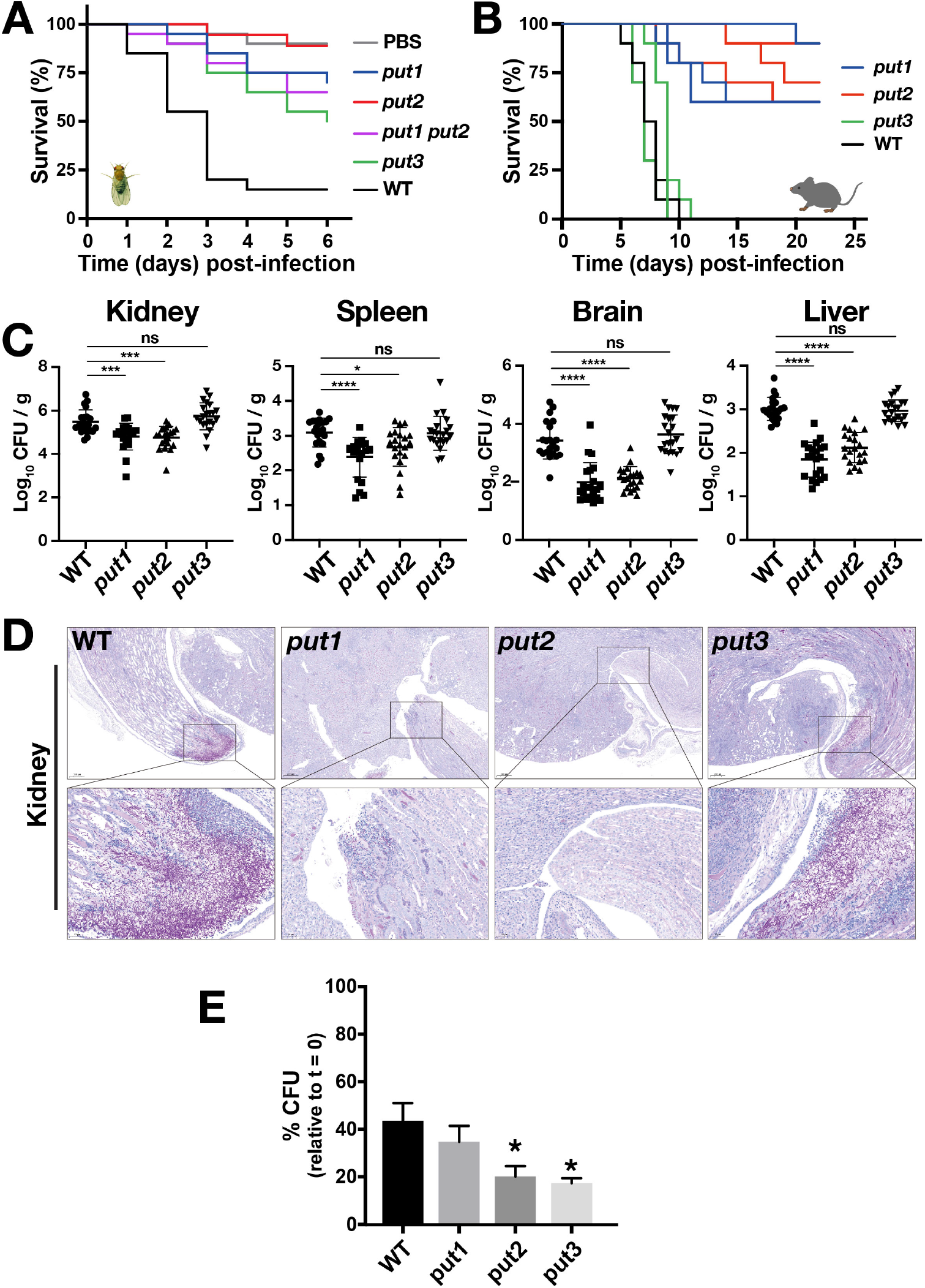
Proline catabolism is required for virulence in *Drosophila* and murine infection models. **(A)** *D. melanogaster Bom*^*Δ55C*^ flies were infected with WT and the indicated *put* mutants and their survival was followed for 6 days. Each curve in the plot were average of at least 6 independent experiments (20 flies/strain), initiated on different days. Strains used: WT (SC5314), *put1* (CFG154), *put2* (CFG318), *put3* (CFG156), and *put1 put2* (CFG159). (\*\*\*\**p*<0.0001 by Log-rank (Mantel-Cox) test). Null strains are designated as e.g., *put1*= *put1-/-*. **(B)** Female C57BL/6 mice were intravenously infected with WT and the indicated *put* mutants and their survival was followed for 23 days. Each curve in the plot were average of 3 independent experiments (10 mice/strain; infected with 2 × 10^5^ CFU/mice). Strains used: WT (SC5314), *put1* (CFG154), *put2* (CFG318), and *put3* (CFG156). **(C)** The fungal load (CFU) in the organs indicated was determined day 5 post-infection. The results from three independent experiments (21mice/strain infected as in B) are plotted. Each symbol represents a sample from an individual mouse (\**p*<0.05,*\*\*\*p<0*.*001*, \*\*\*\**p*<0.0001, one-way ANOVA with Dunnett’s post hoc test). **(D)** Kidneys of infected mice were removed day 5 post infection, fixed in 4% paraformaldehyde at room temperature, sectioned and stained with PAS. Bars = 50 μm. **(E)** Human neutrophils were co-cultured with WT and *put* mutants as indicated. Each bar in the plot is derived from 3 independent experiments with 3 donors with a MOI of 5:1 (*Candid*a:neutrophils). The survival of fungal cells was assessed after 2 h of co-culture. (Ave. ±SEM; \**p* < 0.05 by one-way ANOVA with Dunnett’s post hoc test). Strains used: Wildtype (SC5314), *put1* (CFG154), *put2* (CFG318), and *put3* (CFG156).

Next, C57BL/6 mice were systemically challenged with 10^5^ CFU of wildtype, *put1-/-, put2-/-* or *put3-/-* strains. In comparison to wildtype, *put1-/-* and *put2-/-* strains exhibited significantly attenuated virulence (Fig. 5B). On day 22 post infection, the mean survival rate of mice infected with *put1-/-* or *put2-/-* was 70% and 73.3%, respectively. In striking contrast to the fly model, the *put3-/-* mutant did not exhibit attenuated virulence, which suggests Put3-independent expression of *PUT1* and *PUT2* in mice (Fig. 2D) Consistent to the longer survival times, the fungal burden 5 days after injection were significantly lower in the kidney, spleen, brain, and liver of *put1-/-* and *put2-/-* infected mice (Fig. 5C). Using Periodic Acid-Schiff (PAS)-staining, minimal filamentation was observed in the kidney sections of mice infected with *put1-/-* relative to wildtype, and filamentation was virtually absent in the *put2-/-* strain (Fig. 5D). Interestingly, the kidney infected with *put3-/-* showed smaller areas of infection compared to wildtype, but nonetheless the fungal burden was similar; a result that may be reconciled if the *put3-/-* mutation results in a higher proportion of yeast-like rather than filamentous cells (Fig. 5D).

We recently obtained evidence that proline catabolism is activated in *C. albicans* during co-culture with neutrophils *(46)*. Neutrophils are the most abundant leukocytes in circulation, and play a critical role in controlling and clearing both mucosal and disseminated fungal infections. The regulated generation of ROS within the phagocytic compartment provides the major fungicidal mechanism *(47)*. We examined the survival of *C. albicans* strains when co-cultured with neutrophils. As expected, neutrophils effectively reduced the survival of fungal cells, including wildtype. However, in comparison to wildtype, the *put2-/-* and *put3-/-* mutants exhibited significantly reduced survival. Although, *put1-/-* cells also exhibited reduced survival compared to wildtype, the level of survival was not significantly different (Fig. 5E). Together our results demonstrate that proline is actively assimilated *in situ* and that the capacity to utilize proline is a predictor of virulent growth.

### Visualizing acute *C. albicans* infections of the kidney in real time in a living host

To visualize *Candida*-host interactions we exploited 2-photon intravital microscopy (IVM) to image the early stages of the colonization of mouse kidneys during a systemic infection challenge. Optical access to the kidney was achieved by the implantation of an imaging window. IVM uses lasers to excite molecules in the far-red range, which is more penetrating and less photo-damaging, making it suitable for live tissue imaging. In IVM, only the molecules in the focal volume are excited, thus background signals are very low *(48)*. Furthermore, 2-photon microscopy enables label-free detection of second and third harmonic generation and the intrinsic fluorescence provides information about the tissue morphology and cell metabolism *(49)*. IVM has been successfully applied to study various disease states in live mice, including bacterial infection and cancer *(50)*. IVM provides resolution at the cellular level, and thus has significant advantages over routine *in vivo* imaging methods that rely on bioluminescence to track infections in mice *(51)*.

To observe fungal cells we utilized *C. albicans* strain (PLC096) constitutively expressing a modified version of mCherry (yeast enhanced mRFP, yEmRFP) placed under the control of the strong *ADH1* promoter *(52)*. To monitor the growth characteristics of this strain, we stained yeast-form cells with FITC, which reacts with primary amines on the cell surface, and monitored the induction of filamentous growth (Fig. 6A). Filamentous growth was readily observed, yEmRFP expressing daughter cells emerged from FITC-stained mother cells; FITC remains tightly associated with mother cells. Next, we inactivated *PUT*2 in this strain using CRISPR/Cas9 (CFG479). Using these strains, we proceeded to image the colonization of the kidneys in living BALB/cAnNCrl and specific and opportunistic pathogen free (SOPF) BALB/cByJ mice (Fig. 6B). The SOPF health status proved to be important to minimize individual variation in host responses. An abdominal imaging window was implanted *(53)*, and the kidneys were imaged 4 or 24 h post injection with an inoculum of FITC stained yeast (Fig. 6C). We were able to visualize the renal cortex through the imaging window and reached the superficial areas consisting of proximal and distal convoluting tubules and peritubular arterioles. Sites of colonization were localized with the help of FITC fluorescence, and fungal cell growth and morphogenesis was followed by yEmRFP fluorescence. The autofluorescence of endogenous fluorophores, such as NAD(P)H, and of collagen, were used for navigation and reconstructing tissue morphology. At 4 h post injection, we detected *C. albicans* cells in the renal cortex and in the intertubular space, most likely with capillary or interstitial localization (Fig. 6D). Daughter cells, expressing yEmRFP but lacking FITC-staining, grew forming germ tubes that initiate hyphal growth (Fig. 6D). In addition to IVM, we performed *ex vivo* imaging of acute excised kidneys at 24 h post injection (Fig. 6B, E). The mice injected with wildtype *C. albicans* displayed loci with heavy filamentation with hyphae growing inside and through different tubuli (Fig. 6E). As it is apparent from the autofluorescence, the main source of which is NAD(P)H in the cytoplasm of the tubular epithelial cells, the cellular morphology of the affected tubuli was disintegrated by the growing filaments.

**Fig. 6.**
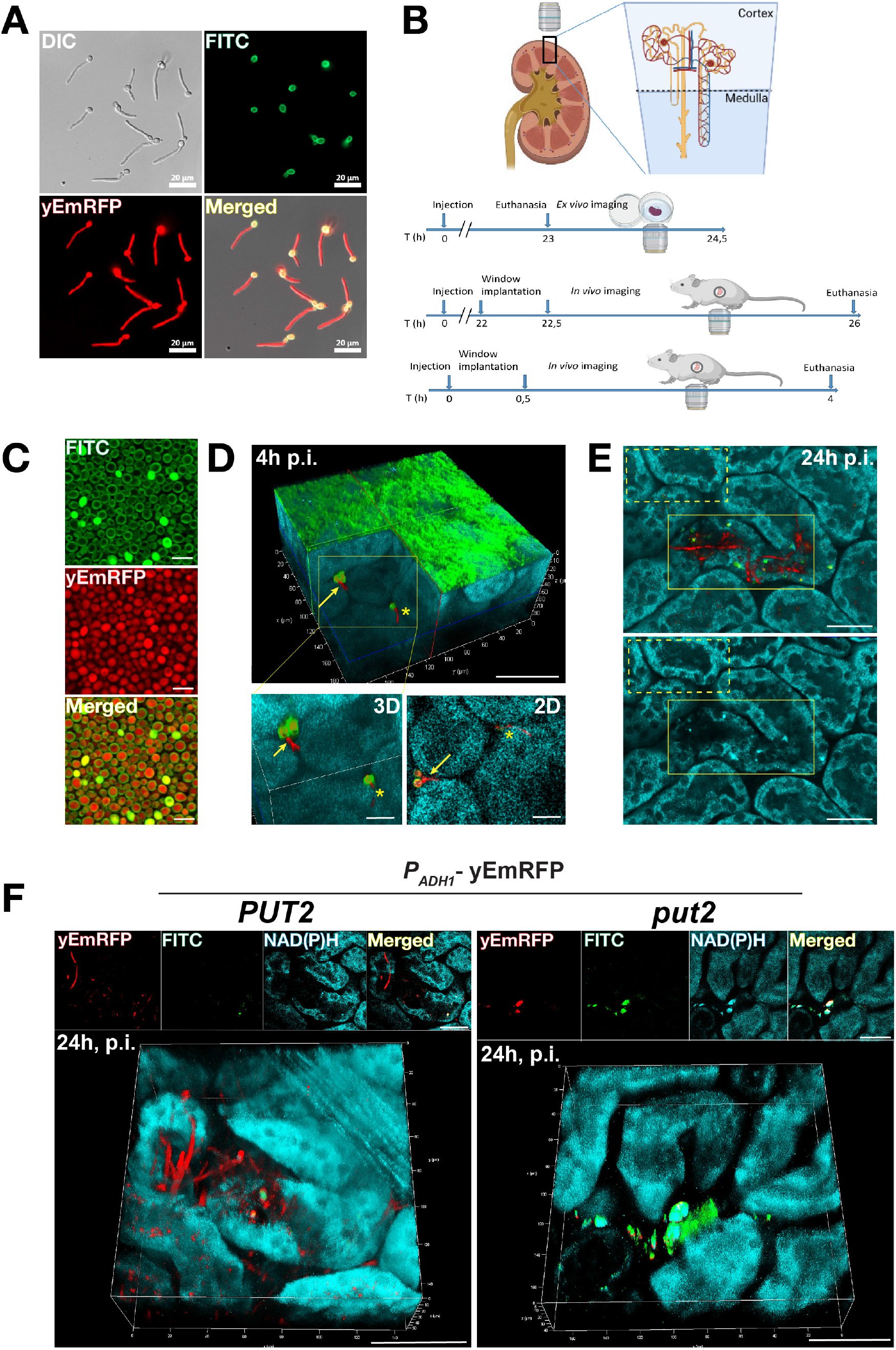
Visualizing acute *C. albicans* infections of the kidney in real time in a living host – Intravital microscopy. (**A**) Exponentially growing *C. albicans* (PLC096) constitutively expressing yEmRFP in YPD were harvested, washed and stained with FITC; under these growth conditions, cells are in budding yeast-forms. The cells were resuspended in DMEM+10% FBS and the induction of hyphal growth was monitored by fluorescence microscopy. Note: FITC fluorescence remains exclusively associated with the mother cells. (**B**) Schematic overview of the imaged kidney, and timelines depicting the experimental design for *ex vivo* imaging 24 h post injection (p.i.), *in vivo* intravital imaging (IVM) between 22.5-26 h p.i., and between 0.5-4 h p.i., respectively. (**C**) Representative aliquot of *C. albicans* (PLC096) cells as in (**A**) prepared for IVM. The micrographs of FITC stained and yEmRFP expressing yeast-form cells were captured with the two-photon microscope used for IVM of infected mice kidneys (panels **D-F**). (**D**) 3D reconstruction of an intravitally acquired z-stack through 90 μm of the renal cortex of a BALB/cAnNCrl mouse injected with *C. albicans* (PLC096). Upper panel, the 3D reconstruction was virtually clipped to expose the intertubular localized *C. albicans* cells. Overlay of 3 channels is shown: yEmRFP (red), FITC (green), autofluorescence of NAD(P)H (teal). Arrow points to a mother cell (yEmRFP and FITC-positive) with an emerging hyphae (yEmRFP positive). The star marks a filament that is penetrating a tubular epithelial cell. The collagen associated with the renal capsule exhibits strong green autofluorescence. Scale bar; 50 μm. Lower panels, higher magnification of the area marked by yellow box in 3D (left) and a representative single 2D plane (right). Scale bars 20 μm. (**E**) Single plane images from a z-stack of *ex-vivo* imaged renal cortex 24 h p.i with *PUT2*+/+ (PLC096). Upper panel, overlay of 3 channels is shown: yEmRFP (red), FITC (green) and autofluorescence of NAD(P)H (teal). yEmRFP and FITC-positive fungal mother cells with long extended hyphal filaments (yEmRFP only) growing through and inside the tubuli. Lower panel, autofluorescence only, allows comparison of the morphologies of renal cells in non-infected (yellow box, dotted line) with infected (yellow box, solid line) areas. Note: the morphology of tubuli in the infected area appear disrupted. Scale bar 40μm. (**F**) 3D reconstructions of intravitally acquired z-stacks through 40-50 μm of the renal cortex of BALB/cByJ SOPF mice, 24 h p.i. with *PUT2*+/+ (*PUT2*, PLC096) and *put2*-/- (*put2*, CFG479) *C. albicans* constitutively expressing yEmRFP. Panels on top of each stack show single 2D plane images for the individual channels and merged. Left panel, *PUT2* mother cells (yEmRFP and FITC positive) with multiple long filaments (yEmRFP positive) are observed to be growing through and inside tubuli. Right panel, *put2* mother cells (yEmRFP and FITC positive) are found accumulated, no filamentation is visible. Scale bar; 50 μm.

To compare the virulence of *C. albicans* wildtype and *put2-/-* strains, we injected two groups of SOPF BALB/cByJ mice. At 24 h post injection, we found foci with filamentous growth in at least 30% of the mice injected with the wildtype strain (Fig. 6F). In contrast, no filamentous growth was observed in mice injected with the *put2-/-* strain. The *put2-/-* cells were found in intertubular locations and were positive for both, yEmRFP and FITC with no sign of filamentation (Fig.6F; see supplementary videos SMov1-SMov4). The lack of filamentation and proliferation of *put2-/-* in the kidney is consistent with the reduced virulence of this strain compared to wildtype in BALB/c mice (Fig. S6) and is comparable to the results obtained in C57BL/6 mice (Fig. 5B).

## Discussion

The results documented here advance our understanding regarding the importance of proline catabolism in fungal infections in four ways. First, many *Candida* spp. known to cause mycoses in humans, including the recently described and multidrug resistant *C. auris*, have the capacity to catabolize proline as sole source of energy. Second, the regulatory mechanisms controlling proline utilization are tightly coupled to mitochondrial function. Third, under respiratory conditions, proline is toxic to cells that lack the ability to catabolize it. This unexpected finding indicates that proline, in addition to the already known toxic intermediate P5C, negatively affects mitochondrial activity. The underlying cause remains to be elucidated, however, it seems likely that when proline accumulates above a critical threshold in mitochondria, it inhibits the activity of a critical ETC component. Fourth, our success in visualizing early events of *C. albicans* cells infecting the kidneys of a living host indicate that single cells respond to *in situ* nutrient cues, which induce invasive filamentous growth in a manner that is dependent on proline catabolism. The knowledge gained and technical advances represented by the successful application of intravital microscopy opens up the new possibilities to address fundamental questions regarding virulent fungal growth with broad biological implications.

Despite proline being one of the first characterized inducers of morphogenic switching in *C. albicans (11)*, we only recently discovered that *C. albicans* can catabolize proline as a preferred energy source *(10)*. The lag in appreciating the physiological significance of proline stems from the over reliance on information obtained in studies with the yeast *S. cerevisiae* (reviewed in *(13)*). Yeast can use proline as a nitrogen source, but is unable to use it as an energy source (Fig. 1). Accumulating evidence indicates that key regulatory differences exist between *C. albicans* and *S. cerevisiae*, which likely reflect the microenvironments in which they evolved (reviewed in *(8)*). *S. cerevisiae* evolved for life in high sugar environments and is widely found in nature. By contrast, *C. albicans* and other members of the *Candida* spp. complex evolved growing in symbiosis with human hosts and are rarely found outside of mammalian hosts. *C. albicans*, unlike *S. cerevisiae*, has a multi-subunit energy conserving NADH dehydrogenase complex (ETC-CI) that couples the oxidation of NADH to ATP production *(54)*. To survive the host environment, fungal cells propagate under similar physiological conditions as human cells. Interestingly, proline catabolism has been implicated as an energy source that facilitates the initiation and progression of multiple invasive diseases, including cancer during nutrient stress *(14)*. Recently, low levels of fungal DNA and cells have been found in association with human cancers *(55)*, suggesting that fungi are able to benefit through spatial association with cancer cells and accompanying macrophages at sites of metastatic growth, perhaps by cross-feeding fungi with host-derived nutrients.

Our finding that multiple *Candida* spp. can catabolize proline as sole source of energy is striking (Fig. 1). Although the PUT pathways are predicted to be conserved among these species, there were clear differences in how well proline is used. We were intrigued to find that the ability to use proline correlated with the known virulence characteristics of the *Candida* species, exemplified by comparing the growth of the two closely related species *C. albicans* and *C. dubliniensis, C. albicans* being more virulent *(56)*. Interestingly, several human pathogens utilize proline to provide energy to facilitate pathogenic growth, including prokaryotes, e.g., *Helicobacter pylori*, and eukaryotes, e.g., *Trypanosomes* and *Cryptococcus neoformans* (reviewed in *(1)*). Future efforts focused on understanding species-specific regulatory differences will be helpful to elucidate the full significance of proline catabolism in pathogenic processes.

The regulation of PUT in *C. albicans* is complex. Put3 expression is subject to glucose repression (Fig. 2C). As Tebung et al. reported *(31)*, proline induces the expression of Put1 and Put2 in *PUT3*-dependent and -independent manner (Fig. 2D). In contrast to Tebung et al., we found that our *put3-/-* strains exhibit residual growth on proline medium (Fig. 1C), which is consistent to the observed levels of Put3-independent expression (Fig. 2D). In the absence of glucose, we found that the expression of cytoplasmic Gdh2 *(8)*, one of the key enzymes in central nitrogen metabolism, is partly regulated by Put3 (Fig. 2D). The inactivation of *PUT3* abrogated both proline- and ornithine-induced Gdh2 expression. *C. albicans*, as a proline prototroph, possesses a full complement of proline biosynthetic genes (*PRO1, PRO2*, and *PRO3*) and hence should be able to convert glutamate to proline. Glutamate, as arginine and ornithine, is metabolized to generate P5C in the cytosol via GSA, which then is reduced to proline by Pro3. We were surprised that in contrast to arginine and ornithine (Fig. 2D and Fig. S2A, C), the presence of exogenous glutamate did not induce Put3-dependent genes. However, the presence of arginine did induce Gdh2 expression independent of Put3, albeit to lower levels, indicating the involvement of transcription factors directly responsive to arginine.

During pathogenic growth of *C. albicans*, the cellular bioenergetic demand requires active mitochondrial function and efficient ATP-generating metabolic processes. The attenuated virulence of *put* mutants, particularly *put2-/-* in both insect and murine infection models (Fig. 5A, B), suggests that proline is indeed actively used in hosts as an energy source. However, our finding that proline at low concentrations (>39 μM) (Fig. 3F) inhibits growth of *C. albicans* cells that lack the ability to catabolize it, complicates this simplistic explanation. In mice, the mean physiological levels of proline, arginine, and ornithine in blood are 269 μM, 137 μM, and 198 μM, respectively *(57)*. In *Drosophila* (larvae), although the level of proline is not markedly high in hemolymph, arginine, which is readily converted to proline in *C. albicans*, is present at high levels *(58)*. These findings also forces us to reconsider why *put* mutants exhibit reduced survival upon phagocytosis by macrophages *(10)*. Phagosomes represent a microenvironment that is thought to be nutrient poor, but clearly has significant levels of proline as judged by robust Put3-dependent expression in phagocytized *C. albicans* cells *(10, 18)*. Consequently, phagocytized *put* mutants are likely to experience proline-dependent inhibition in addition to the inability to catabolize proline. It is noteworthy that in contrast to Put1, Put2 is expressed in the absence of proline and in a Put3-independent manner (Fig. 2D). The level of basal Put2 expression ensures that cells are prepared to oxidize the toxic intermediate P5C when proline becomes available.

Our observations that proline inhibits the growth of *put1-/-* cells albeit to a lower extent than in *put2-/-* cells (Fig. 3A) are consistent with previous reports in yeast (*59*) and plants *(60)*. Presumably proline accumulates in the mitochondria of *put1-/-* where it inhibits an unidentified mitochondrial target. Cells lacking P5C dehydrogenase (*put2-/-*) are substantially more sensitive to proline as a consequence of P5C accumulation (Fig. 3B). Proline as a substrate for ROS formation is supported by several studies using isolated mitochondria, cancer cells and *Drosophila* (reviewed in *(61)*). The high rate of mitochondrial electron transfer associated with proline catabolism may enhance ROS formation. However, such does not appear to be the case in *C. albicans*, as the addition of proline reproducibly decreased ROS in the wildtype cells (Fig. 3C). This may reflect a low reactivity of Put1 towards oxygen, which contrasts with PutA from *Helicobacter pylori* and *H. hepaticus (62)*. Regardless, our data show that ROS is not the primary contributor to proline hypersensitivity of *put2-/-*, as ROS scavengers failed to rescue growth (Fig. 3D and Fig. S3E). Consistent with this notion, it took 6 hours after proline addition before measurable levels of ROS could be reproducibly assayed. It is possible that as *C. albicans* cells detect stress they employ a number of detoxification mechanisms that include superoxide dismutase (SOD) *(63)*, and possibly, the *MPR1*-encoded protein N-acetyltransferase (Mpr1) homologue that detoxifies P5C or GSA by acetylation in yeast *(64)*. We traced the inhibitory effects of P5C to defective mitochondrial respiration (Fig. 3E). Consistently, *put2-/-* mutants are unaffected by proline when grown under mitochondrial repressing conditions in the presence of high glucose (2%) *(10)*. Our findings are well-aligned with Nishimura et al. *(35)*, who showed that strains lacking mitochondrial DNA (*rho*^0^) exhibit diminished P5C hypersensitivity.

The physiological source(s) of proline within the host environment needs to be determined. Although proline assimilation in the host may be a fungal-driven process, where fungal cells actively secrete proteases and cytolytic toxins (candidalysin) *(65)*, it is possible that *C. albicans* cells acquire proline as a result of host-driven processes. There is evidence to suggest that the proteolytic activities in the host, during pathological states such as cancer or sarcopenia contribute to proline availability *(6, 7)*. In addition, other members of the microflora can also facilitate extracellular matrix (ECM) degradation by secreting collagenase during infection, such as been shown in *H. pylori (66)*. In our *in vitro* 3D skin model (Fig. 4E), the source of proline is likely collagen as dermal fibroblasts and keratinocytes secrete matrix metalloproteinases (MMP) *(67)*; the resulting peptide fragments can be internalized by fungal cells and degraded to liberate free proline. Due to its distinct structure, proline induces conformational constraints on the peptide bond, protecting it from degradation by most common proteolytic enzymes *(68)*. Therefore, the release of proline from peptides is thought to have special regulatory and physiological functions. Interestingly, the catalytic release of proline is carried out by a small subset of proteases called prolidase (or proline dipeptidase), which are ubiquitous in nature and are capable of hydrolyzing the bond constrained within the pyrrolidine ring *(68)*. The uncharacterized *C. albicans* ORF (C1_14450C; CGD) encodes a putative prolidase. Supporting the notion that host-driven processes are responsible for proline assimilation, we observed that many proline-rich proteins, including collagen, did not induce the expression of Put1 and Put2 even after 72 h of incubation. Consistently, it took 14 days for the wildtype strain to fully invade the collagen matrix (Fig. 4B), suggesting that collagen is relatively refractory to degradation. In contrast, mucin, an abundant protein in the stomach and mucosal cavities, significantly induced Put1 and Put2, suggesting that *C. albicans* can readily degrade mucin.

IVM enabled unprecedented and unanticipated discoveries of the host-pathogen interactions during early stages of infection of kidneys in a living host. Two-photon microscopy has previously been used to image fungal infections in translucent mouse pinna (ears) *(69)*, however, to our knowledge, our studies represent the first in which *C. albicans* infections have been imaged under physiological *in situ* conditions in a complex intravital organ. The use of *C. albicans* strains constitutively expressing yEmRFP and stained with FITC enabled us to distinguish mother from growing daughter cells. For the first time, we visualized the formation of hyphae in real-time directly deep in the tissue of a living organism (Fig. 6). IVM provided the means to critically examine several assumed, but untested, parameters regarding kidney infections. We found filamentation events in the renal cortex, although these events seem to be rare in the early phase of colonisation. Similar to a bacterial model of infection, where only a few *Escherichia coli* cells were found to be necessary to establish a kidney infection *(50)*, we found that single fungal cells were able to grow and filament (Fig. 6D), suggesting that multiple fungal cells are not required to initiate colonization. We observed that the primary locus for filamentation differed from the sites where non-filamenting *Candida* cells accumulated. Interestingly, filamenting cells localized to the intertubular space, most likely attached to the tubular cells- or capillary wall (Fig. 6D). Another parameter of paramount interest is that we observed *C. albicans* cells transmigrating from the capillary through different endothelial barriers (capillary and tubular) into the lumen of the tubuli (Fig. 6E), thus enabling us to literally observe the establishment of candiduria (urinary tract infections) that typically occurs in patients with systemic candida infections. We also established a connection between *C. albicans* metabolism and virulence using strains defective in proline catabolism that failed to form hyphae under the same *in situ* conditions as the wildtype strain (Fig. 6F).

In summary, the data presented in this work strongly support the idea that proline catabolism is important for *C. albicans* virulence and likely for other *Candida* spp. Our findings indicate that proline is an important energy source when glucose becomes limiting. Importantly, *C. albicans* cells must synchronize the activities of the catabolic enzymes, since proline is toxic, as is the intermediate P5C, making this pathway appropriate to target for the design of antifungal drugs. Small molecules that block Put1 and/or Put2 activity would be expected to inhibit fungal growth. A major outstanding question is how proline is imported from the cytosol to the mitochondria; further studies are warranted to identify a mitochondrial proline transporter. Finally, there is growing evidence that proline serves as an energy source during diverse disease states affecting the host, including cancer and stress, which are risk factors for *Candida* infections. Intriguingly, the expression of proline utilization genes (PRODH) in mammalian cells appears independent of proline, but instead is induced by the p53 tumor suppressor protein, proliferator-activated receptor γ (PPARγ), and/or AMP-activated kinase (AMPK) in response to nutrient and hypoxic stress (reviewed in *(14)*). Given that *Candida* infections are common among immune compromised individuals, proline derived from host-driven degradative processes may be key to understanding fungal virulence. Further work is needed to dissect the complex host-pathogen interactions that impinge on proline catabolism.

## Materials and Methods

### Fungal strains and plasmids

Most of the yeast strains (*Candida* spp. and *S. cerevisiae*) and plasmids (*E. coli*) reported in this work originated or were derived from the Ljungdahl (POL) strain collection (Table S1). *C. albicans* clinical isolates and non-*albicans Candida* species (Table S1) were obtained from several laboratories: Ute Römling (UT), Valerie Diane Valeriano (VDV), Oliver Bader (OB), Matthew Anderson (MA), Steffen Rupp (SR), Constantin F. Urban (CU) and Karl Kuchler (KK). CRISPR/Cas9 plasmids pV1093 and pV1524 were donated by Valmik Vyas (VV). All yeast and *E. coli* strains were stored at -80 °C in YPD with 15% glycerol and recovered as needed on permissive media, and streaked for single colonies.

Fungal strains and plasmids generated in this study are available from the lead contact without restriction.

### Lead contact

Further information and requests for resources should be directed to and will be fulfilled by the lead contact, P.O.L.

### *Drosophila melanogaster* infection model

The *Bom*^*Δ55C*^ *Drosophila* stock was maintained on standard cornmeal agar medium at 25 °C. This fly strain was originally obtained from Bloomington stock center and maintained in Prof. Ylva Engström laboratory, Stockholm University. The *Bom*^*Δ55C*^ mutant flies were collected and transferred to 29 °C for three days prior to injection of fungal cell suspensions.

### Animal studies

All procedures using animals performed at the Experimental Core Facility, Stockholm University, Stockholm, Sweden were approved by the Stockholm Ethics committee (License nr. 9700-2018). Mice were housed in individually ventilated GM500 cages (Tecniplast) under constant humidity (50-60%) and temperature (21 ± 2 °C) and with a constant (year-around) 12-h light/dark cycle. The mice were provided aspen bedding (Tapvei, article no 2212), diet deficient in phytoestrogens (Altromin 1324 variant P, article no 30047), water *ad libitum* and the following enrichment: paper play tunnel (Scanbur, article no 20-CS3B02), aspen gnawing stick (Tapvei, article no 44219999S-brick) and paper nesting material (Scanbur, article no 20-CS1A09). Mice were maintained under Specific Pathogen Free (SPF) conditions according to the FELASA guidelines (2014). For animal experiments performed in Shanghai, China, all procedures were performed in compliance with the protocol approved by the local IACUC committee at the Institute Pasteur of Shanghai, Chinese Academy of Sciences, China (License nr. A2021003) under similar animal care and husbandry conditions as in Stockholm, Sweden. Mice strains and suppliers are listed in Table S1.

### Isolation of human neutrophils

Neutrophils were isolated from the blood of healthy volunteers in compliance with the local ethical committee (Regionala etikprövningsnämnden i Umeå) as approved in permit Dnr 09-210 M with fully informed written consent of donors and all investigations were conducted according to the principles expressed in the Declaration of Helsinki.

### Statistical analysis

Data obtained in this work were analyzed using GraphPad Prism version 9. Specific statistical treatment applied is described in the figure description. In addition, the type of error bars (SEM, SD, or CI (95%)) is dependent on the type of analysis performed. Statistical analysis was performed using the means of at least 3 independent experiments, and statistical significance was determined using unpaired *t*-test, regular one-way analysis of variance (ANOVA) or Kruskal-Wallis (non-parametric) test followed by Dunnett’s multiple comparison test, two-way ANOVA with Sidak’s post hoc test, or Log-rank (Mantel-Cox) test for survival analysis. The following set of notations were used to describe statistical significance: \**p*<0.05, \*\**p*<0.01, \*\*\**p*<0.001, \*\*\*\**p*<0.0001, ns = not significant.

### Exended Method Details

Detailed description of all methods are available in Supplementary Materials under the following subheadings:

**Organisms, media, and culture**

**Genetic manipulation and gene inactivation**

**Reporter strain construction**

**Protein expression analysis**

**Immunoblot**

**Subcellular fractionation**

**Liquid growth assay**

**Assessment of fungal growth on solid media**

**Fungal cell viability assay**

**P5C quantification**

**ROS assay**

**Oxygen consumption measurement**

**ATP quantification**

**Collagen matrix invasion assay**

**Reconstituted human epithelial (skin) (RHE) model**

**Confocal (Airyscan) Microscopy**

***Drosophila* virulence assay**

**Neutrophil killing assay**

**Mouse infection model**

**Intravital and *ex vivo* two (2)-photon microscopy**

## Supporting information

SMov 1. 3D reconstruction of an intravitally acquired z-stack - wildtype C.albicans (Fig 6F, left)

SMov 2. 3D reconstruction of an ex vivo acquired z-stack - wildtype C.albicans

SMov 3. 3D reconstruction of an intravitally acquired z-stack - put2 C.albicans (Fig 6F, right)

SMov 4. 3D reconstruction of an ex vivo acquired z-stack - put2 C.albicans

## Acknowledgments

The authors would like to thank the members of the Per O. Ljungdahl, Claes Andréasson, and Sabrina Büttner laboratories (SU) for their constructive comments throughout the course of this work. Gratitude is extended to the following for supplying strains: Valmik Vyas and Gerald R. Fink (MIT, Cambridge, MA, USA); Valerie Dianne Damiano, Ute Römling (Karolinska Institute, Solna, SE); Måns Ullberg (Karolinska Hospital, Huddinge, SE); Matthew Anderson (Ohio State University, USA); Oliver Bader (University Medical Center Göttingen, Göttingen, DE). We thank Francisco Javier Alvarez and Leopold Ilag for fruitful discussions. We would also like to acknowledge Chris Molenaar of the Imaging Facility-Stockholm University (IFSU), and staff of the Experimental Core Facility-Stockholm University and the Intravital Microscopy Unit-Stockholm University (IVMSU), which is part of the dispersed Swedish National Microscopy Infrastructure (NMI) funded by the Swedish Research Council (VR-RFI 2019-00217).

## Funding

The research in the collaborating laboratories was supported by:

Swedish Research Council 2019-01547, 2022-01190 (POL); 2019-01790 (CP); 2022-00850 (CFU)

Marie Curie - Initial Training Networks (ITN), ImResFun 606786 (POL, KK, SR, TL)

China MOST Key R&D Program 2022YFC2303200; 2020YFA 0907200 (CC)

National Natural Science Foundation of China 32170195 (CC)

Shanghai “Belt and Road” Joint Laboratory Project 22490750200 (CC)

Austrian Science Fund (FWF) ChromFunVir P-32582 (KK)

European Commission, FP7 Framework Programme, Fungitect 602125 (TL)

## Author contributions

Conceptualization: POL, FGSS, MW, CP, BB-V, CC and TJ

Methodology: FGSS, CP, BB-V, CC, TJ, CFU, SR, KK

Investigation: FGSS, CP, BB-V, TJ, AK, KR, NU, SJ, FN and MW

Visualization: FGSS, CP, BB-V, TJ, AK and NU

Funding acquisition: POL, CP, CC, KK, SR, CU and TL

Supervision: POL, FGSS, CP, CC, KK, SR, CFU and TL

Writing – original draft: FGSS, POL

Writing – review & editing: FGSS, POL, CP, CC, SJ and FN

## Data and materials availability

All data are available in the main text or the Supplementary Materials.

## Supplementary Materials

### Extended Method Details (Subheadings)

Organisms, media, and culture

Genetic manipulation and gene inactivation

Reporter strain construction

Protein expression analysis

Immunoblot

Subcellular fractionation

Liquid growth assay

Assessment of fungal growth on solid media

Fungal cell viability assay

P5C quantification

ROS assay

Oxygen consumption measurement

ATP quantification

Collagen matrix invasion assay

Reconstituted human epithelial (skin) (RHE) model

Confocal (Airyscan) Microscopy

*Drosophila* virulence assay

Neutrophil killing assay Mouse infection model

Intravital and *ex vivo* two (2)-photon microscopy

### Figures

**Figure S1. Strain construction and targeted reconstitution**

**Figure S2. Put3-dependent and -independent induction of PUT enzymes**

**Figure S3. Proline catabolic mutants are sensitive to exogenous proline**

**Figure S4. Cells lacking the capacity to catabolize proline exhibit enhanced death**.

**Figure S5. Expression of Proline Utilization enzymes during growth in the presence of different protein sources**

**Figure S6. Mutations inactivating proline catabolism attenuate virulence in BALB/cAnNCrl mice**

### Tables

**Table S1. Key reagents and resources**

**Table S2. Primers used in this study**

### Videos

**SMov 1** Animation of a 3D reconstruction of an intravitally acquired z-stack through the renal cortex of a BALB/cByJ SOPF mouse, 24h p.i. with wildtype *C. albicans*, expressing yEmRFP and stained with FITC, merged channels: yEmRFP (red), FITC (green), Autofluorescence of NAD(P)H (teal) (as in **Fig. 6F, left**).

**SMov 2** Animation of a 3D reconstruction of an *ex vivo* acquired z-stack through the renal cortex of a BALB/cByJ SOPF mouse, 24h p.i. with wildtype *C. albicans*, expressing yEmRFP and stained with FITC, merged channels: yEmRFP (red), FITC (green), Autofluorescence of NAD(P)H (teal).

**SMov 3** Animation of a 3D reconstruction of an intravitally acquired z-stack through the renal cortex of a BALB/cByJ SOPF mouse, 24h p.i. with *put2 C. albicans*, expressing yEmRFP and stained with FITC, merged channels: yEmRFP (red), FITC (green), Autofluorescence of NAD(P)H (teal) (as in **Fig. 6F, right**).

**SMov 4** Animation of a 3D reconstruction of an *ex vivo* acquired z-stack through the renal cortex of a BALB/cByJ SOPF mouse, 24h p.i. with *put2 C. albicans*, expressing yEmRFP and stained with FITC, merged channels: yEmRFP (red), FITC (green), Autofluorescence of NAD(P)H (teal).

