## Supplementary Materials for "Proline catabolism is key to facilitating *Candida albicans* pathogenicity"

Silao *et al.*

Lead author:

#### The PDF file includes:

Extended Method Details (Subheadings):
- Organisms, media, and culture
- Genetic manipulation and gene inactivation
- Reporter strain construction
- Protein expression analysis
- Immunoblot
- Subcellular fractionation
- Liquid growth assay
- Assessment of fungal growth on solid media
- Fungal cell viability assay
- P5C quantification
- ROS assay
- Oxygen consumption measurement
- ATP quantification
- Collagen matrix invasion assay
- Reconstituted human epithelial (skin) (RHE) model
- Confocal (Airyscan) Microscopy
- *Drosophila* virulence assay
- Neutrophil killing assay
- Mouse infection model
- Intravital and *ex vivo* two (2)-photon microscopy

Figs. S1 to S6

Tables S1 and S2

#### Other Supplementary Material for this manuscript includes the following:

Videos:

**SMov1** Animation of a 3D reconstruction of an intravitally acquired z-stack through the renal cortex of a BALB/cByJ SOPF mouse, 24h p.i. with wildtype *C. albicans*, expressing yEmRFP and stained with FITC, merged channels: yEmRFP (red), FITC (green), Autofluorescence of NAD(P)H (teal) (as in **Fig. 6F, left**).

**SMov2** Animation of a 3D reconstruction of an *ex vivo* acquired z-stack through the renal cortex of a BALB/cByJ SOPF mouse, 24h p.i. with wildtype *C. albicans*, expressing yEmRFP and stained with FITC, merged channels: yEmRFP (red), FITC (green), Autofluorescence of NAD(P)H (teal).

**SMov3** Animation of a 3D reconstruction of an intravitally acquired z-stack through the renal cortex of a BALB/cByJ SOPF mouse, 24h p.i. with *put2 C. albicans*, expressing yEmRFP and stained with FITC, merged channels: yEmRFP (red), FITC (green), Autofluorescence of NAD(P)H (teal) (as in **Fig. 6F, right**).

**SMov4** Animation of a 3D reconstruction of an *ex vivo* acquired z-stack through the renal cortex of a BALB/cByJ SOPF mouse, 24h p.i. with *put2 C. albicans*, expressing yEmRFP and stained with FITC, merged channels: yEmRFP (red), FITC (green), Autofluorescence of NAD(P)H (teal).

#### Extended Method Details

##### Organisms, media, and culture

Yeast strains listed in **Table S1. Key reagents and resources** were maintained on solid YPD [1% yeast extract, 2% peptone, 2% glucose (dextrose), and 2% Bacto agar] as a streak from single colonies after recovery from -80 °C glycerol stocks. For yeast selection, YPD is supplemented with nourseothricin (Nou) at the indicated concentrations (200, 100 and 25 µg/ml from a filtered stock solution 200 mg Nou/ml prepared in $H_2O$. Where indicated, 2% glucose in YPD is replaced with 1% glycerol (YPG), 1% lactate (YPL), 0.2% glucose (YPD0.2%), 1% acetate (YPAc), or 2% maltose (YPM). In some specific growth assays, synthetic minimal media were used with the indicated carbon and nitrogen source: SD [0.17% YNB without amino acids and ammonium sulfate (YNB), 38 mM ammonium sulfate (Am, 5 g/L), and 2% glucose], SD* (same as SD but the Am was reduced to 5 mM), SPD (0.17% YNB, 10 mM proline, 2% glucose), SPG (0.17% YNB, 10 mM proline, 1% glycerol), SP (0.17% YNB, 10 mM proline), SGL (0.17% YNB, 10 mM proline, 1% glycerol, 1% lactate). All of the media were prepared from sterile stock solutions prepared in $ddH_2O$ as follows: YP (2X), YNB (1.7%), agar (4%), glucose (40%), glycerol (20%), lactate (20%, pH = 6), sodium acetate (18%), proline (200 mM), arginine (200 mM), ornithine (200 mM), glutamate (200 mM, pH = 6), glutamine (200 mM), and ammonium sulfate (1 M). The pH of lactate and glutamate solutions was titrated using 6M NaOH. All stocks were autoclaved separately except for the amino acids that were filter-sterilized (0.2 µm). Other media modifications or culture conditions are indicated in figure legends or text.

##### Genetic manipulation and gene inactivation

All CRISPR/Cas9 plasmids, repair templates (RT) bearing in-frame stop codons and restriction sites, and verification primers used to generate and verify additional *put* strains were described previously(*1*) and can also be found in **Table S1. Key reagents and resources** and **Table S2. Oligonucleotides**. For plasmids, *E. coli* from glycerol stocks were recovered directly in liquid LB broth with 50 µg/ml carbenicillin and/or 50 µg/ml nourseothricin (Nou) and then processed the following day for plasmid extraction. Specific sgRNAs were cloned in either pV1093 and/or pV1524 backbones(*2, 3*) by blunt-end ligation. Plasmids were verified by sequencing using primer p9. Verified cassettes expressing specific sgRNAs, Cas9, and selection marker, were released from plasmids by *Kpn*I/*Sac*I digestion (~5 µg plasmid/digestion). Digested cassettes were PCR-purified and co-transformed with purified RT (generated by template-less PCR) into the indicated strain using the hybrid lithium acetate/DTT-electroporation method by Reuss, et al.(*4*) with minor modifications described in.(*1*) Nourseothricin-resistant ($Nou^R$) transformants were selected on YPD+200 µg/ml Nou and further clonal purification was made on YPD+100 µg/ml Nou. Purified colonies were verified by colony PCR using the appropriate primers to amplify the mutated gene and then digested with the specific restriction enzyme (*Xho*I) to identify knockouts. For pV1524-derived plasmids, cassette excision was done by growing colonies in liquid YPM and then plated on YPD+25 µg/ml Nou. Nourseothricin sensitive ($Nou^S$) pop-outs were verified by streaking on YPD+100 µg/ml Nou. In most cases, single preparations of purified digested cassettes and RT suffice for 10-15 independent transformations and were kept at -20°C until used. For targeted reconstitution, *put1-/-* and *put2-/-* strains were transformed with wildtype *PUT1* and *PUT2* gene fragments and proline-utilizing ($Put^+$) colonies were selected on SPD resulting in CFG379/CFG380 (*PUT1*+/-) and CFG381/ CFG382 (*PUT2*+/-), respectively.The following primer pairs were used for PCR verification: *put1-/-* (p1/p2), *put2-/-* (p2/p3), *put3-/-* (p5/p6), *PUT1+/-* (p7/p2), *PUT2+/-* (p8/p4).

*C. glabrata PUT1* (CAGL0M04499g) and *PUT2* (CAGL0M04499g) gene deletion were done in the ATCC2001 (CBS138) background strain using the modified SAT1 flipper technique (*4, 5*). Briefly, a YEp352-*NAT1*-Cg*PUT1*urdr and YEp352-*NAT1*-Cg*PUT2*urdr plasmids were generated by Gibson assembly (*6*). Fragments (~10 ng/kb) composed of 500 bp of up- and downstream regions of *PUT1* or *PUT2* coding sequence (CDS), a FRT-FLP-*NAT1*-FRT fragment from pSFS3b (*7*), and an *E. coli* replication origin and an ampicillin resistance marker from YEp352-*SAT1* were assembled using a 2x Gibson assembly master mix (New England Biolabs). The deletion cassettes released by digestion using FastDigest *Nco*I and *Pvu*I, PCR-purified, and transformed into *C. glabrata* by electroporation. Transformants were verified for correct cassette integration and deletion by colony PCR using DreamTaq Green DNA Polymerase.

##### Reporter strain construction

The triple tagged *C. albicans* reporter strain co-expressing Gdh2-GFP, Put1-RFP, and Put2-HA was constructed via a series of homologous recombination transformations (Fig. S1D) with specific cassettes described previously(*8*). The GFP tag uses *URA3* as marker, while both the RFP and HA tagging cassettes that were derivatives of the SAT1-flipper cassette (*4*) use $Nou^R$ as a recyclable resistance marker. Starting from strain CFG407, which already co-expresses Gdh2-GFP and Put1-RFP, this strain was subjected to another round of transformation with a ~4.8 kb *PUT2-HA-CaSAT1* cassette amplified from pFA6a-3xHA-SAT1-FLP using primers p10/p11 generating the $Nou^R$ strain CFG430. Cassette excision was made by growing CFG430 in YPM generating the $Nou^S$ pop-outs (CFG433). To increase the flexibility of this strain for western blot works, we created strain CFG438 by transforming CFG433 with pJA21 digested with *Kpn*I/*Sac*I, which releases the $P_{ADH1}$*-RFP-caSAT1* fragment, enabling CFG438 to constitutively express free RFP. CFG438 was popped-out to excise the cassette generating the $Nou^S$ strain CFG441. For the creation of Put2-GFP strain expressing free RFP,

CFG219 (*PUT2/PUT2-GFP-URA3*) strain was transformed with *Kpn*I/*Sac*I-digested pJA21 to generate CFG237 ($Nou^R$) and then popped-out to obtain CFG259 ($Nou^S$). Strains CFG441 and CFG259 were used as backgrounds for CRISPR/Cas9-mediated inactivation of *PUT3* using plasmid pFS084 resulting to strains CFG443 and CFG301, respectively. For the creation of Put3-HA expressing strains, SC5314 was transformed with the *PUT3-HA-CaSAT1* cassette amplified from pFA6a-3xHA-SAT1-FLP using primers p19/p20, which then generate the strains CFG187/CFG188 (independent clones from two separate transformations).

PCR conditions for cassette amplification using either Ex-TAQ polymerase (Takara) or Phusion polymerase (Thermo Fischer Scientific) in a 300 µl (50 µl x 6) reaction volume are as follow: 1) Initial melting: 98°C, 30s; 2) 3-step: Melting: 98°C, 10s; Annealing: 55°C, 30s; Extension: 72°C, 5 min; 3) 2-step: Melting: 98°C, 10s; Extension: 68°C, 5 min; 4) Final extension: 68°C, 10 min; and 5) 4°C. After amplification, samples were pooled and then an aliquot run on 1% agarose to assess successful cassette amplification. Pooled PCR samples were purified and then stored at -20°C until used. Purified cassettes (0.2-0.5 µg) were transformed into the indicated strains and then the correct clones were screened by colony PCR, complemented by immunoblotting and/or fluorescence microscopy. In most cases, a single preparation of purified tagging cassettes is good for 10-15 independent transformations and are kept at -20°C until used. The following primer pairs were routinely used for PCR verification of cassette integration: *PUT1-RFP-CaSAT1* (p14/p16), *PUT2-HA-CaSAT1* (p12/p13), *GDH2-GFP-URA3* (p17/p18), $P_{ADH1}$*-RFP-CaSAT1* (p15/p16), *PUT3-HA-CaSAT1* (p5/p13).

#### Protein expression analysis

To analyze the expression of PUT enzymes in the presence of different nitrogen sources, overnight SGL cultures of the indicated strains were diluted into 3 ml of fresh SGL medium at $OD_{600} \approx 0.3$. Cultures were grown to log phase with vigorous shaking for 6 h at 30 °C and then a ~0.8 ml aliquot was added into an Eppendorf tube containing the required volume of proline (or other nitrogen source) to achieve the required 10 mM inducing concentration. Induction was performed on a table-top thermo-shaker (900 rpm, 30 °C) for 1 h. After induction, whole cell lysate was prepared by NaOH/trichloroacetic acid (TCA) precipitation; briefly, cells were mixed with 280 µl of ice-cold 2 M NaOH with 7% β-mercaptoethanol (βME) to the tube and then incubated for 15 min on ice. Lysed cells were mixed with an equal volume of cold 50% TCA to precipitate proteins overnight at 4°C. Protein precipitates were collected by centrifugation at 17,000*g* (4°C, 10 min), solubilized completely in the required volume of 2X SDS sample buffer containing 5% βME and 167 mM of Tris, and then boiled at 95-100°C for 5 min. Samples were either processed immediately for western blot or stored at -20°C until run. For Put3 expression analysis, strains (CFG187/CFG188) expressing Put3-HA were first grown overnight in either YPD or SGL. The following day, cultures were diluted to $OD_{600} \approx 0.3$ in their respective medium where they are allowed to grow exponentially for 4 h (YPD) and 6 h (SGL) at 30 °C before collecting cells for whole cell lysate preparation by NaOH/TCA method and western blot.

For induction of PUT enzymes during growth on protein as sole nitrogen source, strain CFG433 was grown in modified Hanks Balanced Salt Solution (HBSS) [NaCl (138 mM), KCl (5.33 mM), $KH_2PO_4$ (0.44 mM), $Na_2HPO_4$ (0.3 mM), $MgCl_2$ (0.5 mM), $MgSO_4$ (0.41 mM), $CaCl_2$ (1.26 mM), $NaHCO_3$ (4 mM), HEPES (25 mM, pH = 7.4), Biotin (2 µg/ml), 3.8 mM glucose, and 0.83 mM lactate] containing the following nitrogen sources: collagen (from human placenta), human serum albumin, hemoglobin (from bovine blood), mucin (from porcine stomach Type II) or 10 mM ammonium sulfate. These proteins were aseptically dissolved in HBSS at 0.5 mg/ml prepared fresh for every experiment. This HBSS was modified to reflect the level of glucose and lactate in interstitial fluid (*9*). To maintain sterility, this medium was supplemented with Penicillin and Streptomycin (Penn/Strep) commonly used in cell cultures; 100-200 µl aliquots of these solutions were spotted on YPD for sterility check. We used collagen from human placenta (Type IV) since it is readily soluble in this buffer. Other types of collagen (e.g., from bovine Achilles tendon or rat tail) were inappropriate for direct comparison since they require acetic acid for dissolution. To start the experiment, a single CFG433 colony from freshly streaked YPD plates were pre-cultured in 3 ml of SGL medium for 8 h and then afterwards diluted in 50 ml of fresh SGL at $OD_{600} \approx 0.05$ followed by growth for 16 h at 30 °C. Cells were washed twice with $ddH_2O$ and then resuspended in 0.5 ml of $ddH_2O$ to concentrate the cells. Washed cells were added to 3 ml of medium in a 6-well microplate at $OD_{600} \approx 1$. Plates were incubated static in a humidified incubator at 37 °C with 5% $CO_2$ for 1 or 3 days. After the indicated timepoint, cells were recovered gently using rubber scraper and transferred into a 15 ml falcon tube. The wells were washed twice with 3 ml of ice cold $ddH_2O$ and the washings pooled into the same 15 ml tube. The cells were washed twice and resuspended in 1 ml of cold $ddH_2O$. A 0.8 ml aliquot of this is subjected to whole cell lysis by NaOH/TCA method and the rest analyzed by $OD_{600}$. Protein pellets were solubilized in appropriate volume of 2X SDS sample buffer calculated from $OD_{600}$, boiled, and then subjected to immunoblotting to detect the indicated proteins.

#### Immunoblot

Denatured proteins were first resolved in 4-12% Bis-Tris pre-cast gel in 1X NuPAGE MOPS SDS running buffer (150V, ~1.5 h) and then (electro)transferred onto a nitrocellulose membrane for 1 h in a 1X Tris-glycine transfer buffer with 20%

ethanol (120V, ~1.5 h) in a Bio-Rad Mini Trans-Blot® Cell apparatus. After protein transfer, membranes were blocked with gentle agitation in 10% skimmed milk (MSK) solution in TBST for 1 h. Target proteins in the membrane were probed either individually or simultaneously using an optimized antibody cocktail prepared in 5% MSK/TBST solution; primary antibody cocktail (overnight incubation at 4°C): α-GFP (1:3000), α-mCherry (1:6000), α-Tdh3 (1:5000), α-actin (1:5000) and secondary antibody cocktail (1 h incubation at room temperature): goat α-mouse poly-HRP (1:15000), goat α-rabbit poly-HRP (1:10000), α-HA-HRP (1:15000), α-tubulin-HRP (1:2500). After extensive washing with TBST, membranes were added with chemiluminescent substrate (SuperSignal West Dura Extended Duration Substrate) to detect the immunoreactive bands using the Azure 280 or ImageQuant LAS 500 detection system. Signal quantification on TIFF files were made using Image Lab (Bio-Rad).

**Subcellular fractionation**

For subcellular fractionation analysis, *C. albicans* cells (CFG433) were collected from overnight YPD culture (100 ml), washed twice with $ddH_2O$, and then inoculated into 100 ml of prewarmed YPG (in 500 ml flask) at $OD_{600} \approx 2$. Cultures were grown under aeration (>150 rpm) for 4 h at 30 °C before harvesting cells for cytosolic and mitochondrial fractionation according to the protocol by Meisinger et al. (2006)(*10*) with minor modifications described in our previous work (*8*). Briefly, spheroplasts prepared using Zymolyase-100T were homogenized in a glass-Teflon homogenizer and then the diluted lysate (L) was subjected to two low-speed centrifugation steps to remove cell debris and unbroken cells. The supernatant obtained after the second centrifugation step (4,000*g*, 5 min, 4°C; SL 40R, Thermo Scientific) was labeled as the total cell lysate (T) and subjected further to high-speed centrifugation at 12000*g* (JA-14.50, Beckman Coulter) for 15 min at 4°C. The resulting crude mitochondrial (pellet) and cytosolic (supernatant) fractions were further processed to remove contaminating components from either the cytosol or mitochondria, respectively (See Fig. 1B for scheme). One (1) ml of crude cytosolic fraction was spun down on a table top centrifuge at 17,000*g* for 10 min at 4°C and the resulting supernatant (C) was used. To further purify the crude mitochondrial fraction, the pellet was washed twice without resuspension with 20 ml of cold homogenization buffer and then the pellet resuspended gently in 1 ml of ice-cold SEM buffer; the mitochondrial suspension was first centrifuged at low speed (4000*g* for 10 min) and the resulting supernatant was centrifuged at 12000*g* for 10 min to collect the pellet highly enriched in mitochondria (M). Protein suspensions with normalized concentration (20 mg/ml) were diluted in 2X SDS sample buffer and then processed for immunoblot (see Western blot section) but with the following antibody cocktail: primary: α-GFP (1:3000), α-mCherry (1:6000), α-Tdh3 (1:5000), α-ATP5a (1:2000), and secondary: goat α-mouse poly-HRP (1:15000), goat α-rabbit poly-HRP (1:15000), α-HA-HRP (1:15000).

**Liquid growth assay**

For growth in liquid synthetic minimal medium in tubes, the indicated strains picked from single colonies were pre-cultured overnight in SD or SGL at 30 °C with aeration. The following day, cultures were refreshed in the same medium at a starting $OD_{600} \approx 0.3$ and volume of 6 ml. After 3 h of growth, cultures were split equally into two tubes (3 ml/tube) and then added with either proline (10 mM) or equal volume of $ddH_2O$. Spiked cultures were incubated for 20 h prior to $OD_{600}$ measurement using portable cell density meter (WPA CO 8000, Biochrom, UK). For growth with antioxidants (N-acetyl cysteine (NAC), Mito-TEMPO, or TIRON), cells were grown for 2 h before splitting the cultures (3 ml/tube). These cultures were diluted 1:1 with 3 ml of fresh medium without or with 2X strength of the compound (dissolved or added directly in the medium followed by 0.2 µm filter-sterilization). Cultures were incubated for another 1 h before spiking with proline (10 mM) or $ddH_2O$ and the growth determined 20 h later. For growth in rich complex medium, cells from overnight YPD cultures were washed at least 2X with $ddH_2O$ and then inoculated into 4 ml of YPD, YPG, YPL, and YPAc at a starting $OD_{600} \approx 0.01$ after which growth was recorded 24 h later. When flocculation is to be observed, cultures at the indicated timepoints were vortexed and allowed to stand for at least 3 min prior to photograph. For high throughput growth assays in microplate format, cells of the indicated genotype pre-grown in SGL were inoculated directly into 10 ml of SGL medium at $OD_{600} \approx 0.05$. Using a multi-channel pipette, 100 µl aliquots of this adjusted cell suspension were transferred into 96 well plate followed by the addition of the indicated concentrations of proline or $ddH_2O$. Growth (as absorbance at 600 nm) was monitored every 5 min in a TECAN microplate reader set at 30 °C with continuous shaking.

**Assessment of fungal growth on solid media**

To assay growth on solid medium by drop plate, cells were collected from overnight YPD or SD cultures, washed twice in $ddH_2O$, and then adjusted to $OD_{600} \approx 1$. Five microliters of 10-fold serially diluted cell suspension in $ddH_2O$ prepared in sterile 96-well microplate were spotted onto the solid medium. Spots were allowed to dry before incubating the plates inverted at 30 °C for 48 h (2 days) and photographed. Filamentation on solid Spider medium was carried out as previously described(*8*). Briefly, cells from overnight YPD cultures were collected, washed three times in PBS, and then a 5 µl aliquot of $OD_{600} \approx 1$ cells was spotted on the surface of Spider medium (1% nutrient broth, 1% mannitol, 0.2% $K_2HPO_4$, 1.35%

agar, pH = 7.2) first reported by Liu *et al.*(*11*). Macrocolonies were examined and photographed after 6 days at 37 °C. Other specific growth conditions are indicated elsewhere in the text.

**Fungal cell viability assay**

To assay cell death on macrocolonies, Phloxine B (PXB) was used as a viability stain(*12*); PXB accumulates in dead cells that lack the metabolic energy to extrude it. Briefly, cells from overnight YPD cultures were harvested, washed twice in dd$H_2O$, adjusted to $OD_{600} \approx 1$, and then a 5 µl aliquot was spotted on the indicated solid medium with 10 µg/ml of Phloxine B followed by incubation at 30- or 37 °C for 3 days. PXB plates were prepared by aseptically adding the appropriate volume of PXB (10 mg/ml in dd$H_2O$ stock) to molten agar (~75-85°C) and held in this temperature for several minutes (>10 min) to ensure sterility. For cells grown in liquid cultures, propidium iodide (PI) was used as a viability stain; PI is a membrane impermeant dye that is excluded from viable cells and that intercalates to DNA of dead necrotic cells to exhibit bright red fluorescence. Briefly, cells were grown as indicated in the specific text and then a 1 ml aliquot of the culture was mixed with 1 µl of PI (1 mg/ml in $H_2O$) to a final concentration of 1 µg/ml. Within 3 min after adding the dye, the cells were examined under fluorescence microscope (Cell Observer or Axiocam Observer 7) in the red and DIC channels. Images were captured from at least 3 different frames.

**P5C quantification**

From a 30 ml log phase SGL culture ($OD_{600} \approx 1.5 – 2.5$) of each indicated strain, 6 ml aliquot were transferred to separate tubes containing proline (10 mM final concentration) or dd$H_2O$. Cultures were incubated for 2 h at 30 °C with aeration and then a 100 µl aliquot was transferred to each well of a round bottom 96-well microplate placed on ice. Exactly 100 µl of 10% TCA was added to each well and were allowed to incubate for 15 min protected from light. The direct extraction of P5C in TCA is crucial as P5C is stable in acidic condition (*13*). Using a multichannel pipette, a 50 µl volume of 2-aminobenzaldehyde (2-ABZ) solution in 99.5% ethanol was added to each well and then incubated on ice for another 5 min. Plates were centrifuged at room temperature for 5 min at 4,000 rpm (SL 40R, Thermo Scientific). Supernatants (100 µl) were transferred to a fresh 96-well microplate (clear, flat bottom) and then immediately analyzed for absorbance at 444 nm using the Enspire microplate reader (Bio-Rad). All readings were normalized to cell density ($OD_{600}$) and were presented as fold change relative to wildtype strain grown without proline.

**ROS assay**

For detection of total ROS in strains grown in the presence or absence of exogenous proline, the chemiluminescent luminol-horse radish peroxidase (HRP) system was used. Luminol (Cat. # 123072, Sigma-Aldrich) was dissolved in DMSO (200 mM stock) while HRP (Type 1; Cat. # P8125, Sigma-Aldrich) was dissolved in PBS (300 units/ml stock). From a 30 ml log phase SGL culture ($OD_{600} \approx 1.5 - 2.5$), a 6 ml aliquot was spiked with either 10 mM proline (final concentration) or equal volume of dd$H_2O$. Cultures were grown for 6 h at 30 °C and then a 100 µl aliquot was transferred to each well of a flat-bottom Nunc 96-well white microplate. For control samples, $H_2O_2$ (50 mM; positive control), N-acetyl cysteine (NAC, 10 mM; general ROS scavenger), MitoTempo (MT, 100 µM; mitochondrial-specific superoxide scavenger) was added to the indicated tubes 30 min (i.e., after 5.5 h) before transferring to microplate. Using a multi-channel pipette, a 100 µl of 2X strength luminol-HRP mixture in 100 mM HEPES (pH = 7.4) was added to each well and then mixed; the final concentration of these components per well are as follows: luminol (200 µM), HRP (0.12 units), and HEPES (50 mM). Plates were kept at room temperature for 10 min protected from light before reading the luminescence in Enspire microplate reader (Biorad). Signals were acquired in plate mode set at 30 °C, 1 sec integration time and repeated every 3 min for at least 45 min. All readings were normalized to cell density ($OD_{600}$) and results were presented as both signal accumulation over time and area under the curve (AUC).

**Oxygen consumption**

Cells of the indicated strains were first grown aerobically to log phase ($OD_{600} \approx 1.5 - 2.5$) in a baffled flask containing 20 ml of SGL. From this culture, 4 ml were added to tubes containing proline (10 mM final concentration) or dd$H_2O$. Cultures were incubated for 4 h in a shaking incubator (30 °C) to allow for enzyme expression and then analyzed immediately for oxygen consumption. Oxygen consumptions were measured at 30 °C using a high-resolution oxygraphic system (Oxytherm+R system, Hansatech Instruments Ltd) calibrated using air-saturated dd$H_2O$ and sodium sulfite ($Na_2SO3$) for maximum and zero oxygen levels, respectively. Data acquisition was performed using OxyTrace+ Windows® software every 1 sec. Briefly, 1.7 ml of pre-warmed (30 °C) cellular respiration buffer (1xPBS supplemented with 407 µM $MgSO_4$, 493 µM $MgCl_2$, 1.26 mM $CaCl_2$, 0.2% BSA) was added into the oxygen electrode chamber and then the signal allowed to stabilize for ~3 min with stirrer speed set at 75 rpm. Using a 1 ml syringe with long needle (Sterican; 23G x 2 3/8", Ø0.60 x 60 mm), a 0.3 ml aliquot of the culture is added into the oxygen electrode chamber via the injection port and then the signal acquired for ~5 min. While reading, an aliquot of the culture is also diluted for viable cell count. Rates (in nmol $ml^{-1}$ $min^{-1}$) were obtained from the slopes (3 min) derived using the line of best fit function and then normalized to cell density.

**ATP quantification**

For measurement of intracellular ATP content, a bioluminescence-based detection kit (#A22066; Molecular Probes, Invitrogen) was used. Colonies of the indicated strains from freshly streaked glycerol stocks were resuspended in 300 µl of $ddH_2O$ and diluted in 6 ml of fresh YPD at $OD_{600} \approx 0.01$ followed by growth for 16 h at 30 °C. Cells were then harvested, washed twice with sterile ice-cold Tris Buffered Saline (TBS; 50 mM Tris-HCl, pH 7.5, 150 mM NaCl), and then resuspended in TBS. Cell density was adjusted to $OD_{600} \approx 20$ in a 1 ml total volume. Cells were then harvested at 10,000*g* for 3 min (4 °C) before re-suspending the entire pellet in a modified TCA buffer containing 100 mM Tris-HCl (pH = 8.0), 2% trichloroacetic acid (TCA), 25 mM ammonium acetate, and 4 mM EDTA. TCA concentration was reduced to 2% from the previously used concentration of 10% (*1*) to achieve a final concentration of < 0.01% TCA following dilution in the luciferase reaction. Cell suspension was transferred to pre-chilled tubes containing glass beads and then subjected to bead beating (Bio-Spec; 5 × 1 min, 4 M/s with 2 min on ice between pulses). Cell lysates were collected and a portion of the supernatant was first diluted 20X using Tricine buffer (pH = 7.5) and then analyzed for ATP following the manufacturer's instruction. Luminescence was analyzed using microplate reader (Berthold) with 1 sec integration time. A portion (400 µl) of the same lysate was used to determine total protein concentration. Briefly, the lysate was mixed with 77 µl of 50% TCA to bring back the TCA concentration to 10%. After 15 min incubation on ice, precipitated proteins were collected by centrifugation at max speed (17000*g*) for 10 min (4 °C). Supernatant was discarded and then the pellet mixed with 30 µl of 0.2 M NaOH. The neutralized pellet was dissolved further in 370 µl of RIPA buffer (pH = 8.0) to resolubilize the protein. After lysate clarification at 12, 000 rpm for 10 min (4 °C) the supernatant was diluted and then analyzed for protein using the bicinchoninic acid (BCA; Sigma) assay. Results presented are average of ATP normalized to total protein concentration analyzed from 4-5 biological replicates performed in duplicate.

**Collagen matrix invasion assay**

Prior to start of the assay, a 400-µl aliquot of ice-cold PureCol™ EZ gel (Cat. # 5074; Sigma-Aldrich) was added onto each transwell (PET membrane, pore size = 8 µm; VWR International, PA, USA) in a 24-well microplate and then allowed to solidify for 1 h at 37 °C in a humidified $CO_2$ incubator. For fungal cell preparation, cells from log phase YPD culture were harvested, washed twice in PBS, and then diluted to $OD_{600} \approx 0.1$. A 1 µl aliquot of the adjusted cell suspension was carefully added directly on top of the collagen. Each transwell was aseptically transferred to a separate well containing 1 ml of complete D10 medium acting as the receiving medium. For competition assay, cell suspension of both wildtype and the *put* mutant were mixed 1:1 and then a 1 µl aliquot was added directly on top of the matrix. Fungal cells were allowed to invade the matrix for 14 days in a humidified chamber set at 37 °C with 5% $CO_2$. In each assay, the *cph1*Δ/Δ *efg1*Δ/Δ strain (CASJ041)(*14*) was used as non-filamenting control. For assessment of individual strains, the receiving medium was serially diluted and CFU determined. For competition assay, at least 50 colonies on plates from appropriate dilutions were scored for growth on SPD plate to determine the wildtype or *put* mutants (i.e., growth = wildtype; no growth = *put* mutant) and were compared against the input ratio. For wildtype or *cph1*Δ/Δ *efg1*Δ/Δ scoring, colonies were spotted on Spider medium and grown for up to 6 days at 37 °C where the wildtype strain undergoes heavy wrinkling due to filamentous growth. Results presented are from 5-7 biological replicates.

**Reconstituted human epithelial (skin) (RHE) model**

For the generation and infection of human in vitro skin model derived from S1F (immortalized human dermal fibroblasts) and Ker-CT (immortalized human keratinocytes) cell lines, the procedure was performed essentially as described (*15*) with minor modifications with respect to media exchange to keep the glucose levels down enabling relief from glucose repression in *C. albi*cans. In this specific work, media replacement, which is usually performed directly prior to infection, was halted 3 days prior to infection.

**Confocal (Airyscan) Microscopy**

For microscopic analysis, strain (CFG407) expressing both Gdh2-GFP and Put1-RFP was grown as in for subcellular fractionation i.e., 4 h growth in YPG at 30 °C. Cells were collected, washed, and then resuspended in PBS before viewing the cells using laser scanning confocal microscope (LSM800, 63x oil; Airyscan mode) in the green and red channels excited with 488 nm and 561 nm lasers, respectively, with Differential Interference Contrast (DIC) taken separately. Z-stack confocal images were obtained and the raw Airyscan images were processed using the Zen Blue software's Airyscan processing function.

##### *Drosophila* virulence assay

Infection of *D. melanogaster Bom*$^{\Delta 55C}$ flies was performed as described (*16*). Flies were injected with approximately 500 cells/fly and then maintained in separate vials where fly survival was monitored for at least six days at 29 °C.

##### Neutrophil killing assay

To assess the sensitivity of the proline catabolic mutants towards neutrophil attack, fungal cells were co-cultured with human neutrophils freshly isolated from peripheral blood of healthy human donors using Histopaque 1119 (Sigma-Aldrich) described in (*17*). Neutrophils in RPMI1640 medium without phenol red were seeded at 100,000 cells/well (24-well plate) and then incubated in a humidified chamber (37 °C, 5% $CO_2$) for at least 30 min to allow the neutrophils to adhere to the culture dish. For the preparation of fungal inocula, *C. albicans* cells were first grown overnight in YPD, subcultured to log phase, washed, and then adjusted to ~$10^6$ CFU/ml in RPMI. *C. albicans* cells were mixed with neutrophils at MOI of 5:1 (*Candida*:neutrophils) and then co-cultured for 2 h (t = 2) in the humidified chamber after which the neutrophils were lysed to release fungal cells. Lysates were diluted and plated on YPD agar to determine viable fungal count. For t = 0, neutrophils were immediately lysed after adding the fungal cells. The % killing after t = 2 was calculated based on the CFU obtained initially at t = 0.

##### Mouse infection model

For the murine model of hematogenous disseminated candidiasis, the wildtype *C. albicans* SC5314 and the three *put* mutants were individually inoculated to YPD broth and grown overnight at 30 °C. Cells were harvested and washed three times with phosphate-buffered saline (PBS), and counted using hemocytometer. A group of 6–8-week female C57BL/6 mice (n=10) were intravenously inoculated with each of the indicated strains, using an inoculum of 2x$10^5$ CFU per mice. The mice were monitored once daily for weight loss, disease severity and survival. The survival curves were statistically analyzed by the Kaplan-Meier method (a log-rank test, GraphPad Prism). The fungal burden was assessed by counting CFU. Mice were euthanized on the 5$^{th}$ day after infection. The organ tissues, including kidneys, spleen, brain and liver, were collected, homogenized and appropriately diluted for plating on Sabouraud Dextrose Agar (SDA) medium. After incubation at 30 °C for 48 h, colony counts were performed and data were statistically analyzed. For histological analyses, the mice were euthanized on the 5$^{th}$ day after infection and the kidneys were fixed in 4% paraformaldehyde at room temperature. The sections were stained with periodic acid-schiff (PAS) and photographed.

For systemic infection in BALB/cAnNCrl mice performed in Stockholm, a group of at least ten 6- to 8-week-old mice per strain were infected via the lateral tail vain with 5x$10^5$ CFU of *C. albicans* wildtype (SC5314) or *put2*-/- (CFG318) mutants. Survival was scored daily; moribund mice were sacrificed and death was recorded the following day. For assessment of fungal burden in the kidney, mice from each group were sacrificed (2 days) and kidneys removed and homogenized in PBS by bead-beating using glass plating beads (4.5 mm). CFU counts were determined from diluted tissue homogenates plated on YPD agar with 30 µg/ml chloramphenicol. Survival curves between wildtype and *put2*-infected mice were analyzed by the Kaplan-Meier method whereas the difference in fungal burden was analyzed by student *t*-test, both performed in GraphPad Prism.

##### Intravital and *ex vivo* two (2)-photon microscopy

BALB/cAnNCrl and BALB/cByJ SOPF mice (8-10 weeks old) were injected into the tail vein with 5x$10^5$ CFU of *C. albicans* wildtype or *put2-/-* (both constitutively expressing a modified version of mCherry (yEmRFP) under the control of the *ADH1* promoter) in sterile physiological saline. For *ex vivo* imaging, the animals were sacrificed 24h post injection, the kidneys were removed, submerged in physiological saline and imaged immediately for max. 90 min. For intravital imaging, the mice were – immediately or 22-24h post injection- anesthetized with isoflurane, implanted with an abdominal imaging window above the left kidney, transferred immediately to the microscope and imaged for a maximum of 4h under isoflurane anesthesia, a heating blanket and monitoring of body temperature. 100µl/h physiological saline was injected subcutaneously to prevent dehydration. After the last image acquisition, the animals were euthanized by cervical dislocation. Intravital and *ex vivo* microscopy was performed using a Leica SP8 DIVE inverted confocal laser scanning microscope (Leica Microsystems, Wetzlar, Germany), equipped with a pulsed multiphoton laser INSIGHT DUAL X3 with 80MHz repetition rate (Spectra Physics, MKS Instruments, Inc., Andover, Massachusetts, USA) and 4Tune spectral non-descanned detectors. Excitation occurred with a laser power of 5-8% and at 1075nm (mCherry), at 930nm (FITC) and at 780nm (autofluorescence), fluorescence was collected at 580-680nm (mCherry), at 492-550nm (FITC) and at 380-460nm (autofluorescence). Images were acquired with HC PL APO 20x/0.75 CS2 objective lenses (Leica Microsystems, Wetzlar, Germany). Image acquisition, deconvolution and data analysis was performed using the software platform LAS X and integrated Hyvolution function (Leica microsystems, Wetzlar, Germany), enabling deconvolution of with Hygens Essential (Scientific Volume Imaging, Hilversum, Netherlands). The cartoons were created with the help of Biorender.com.

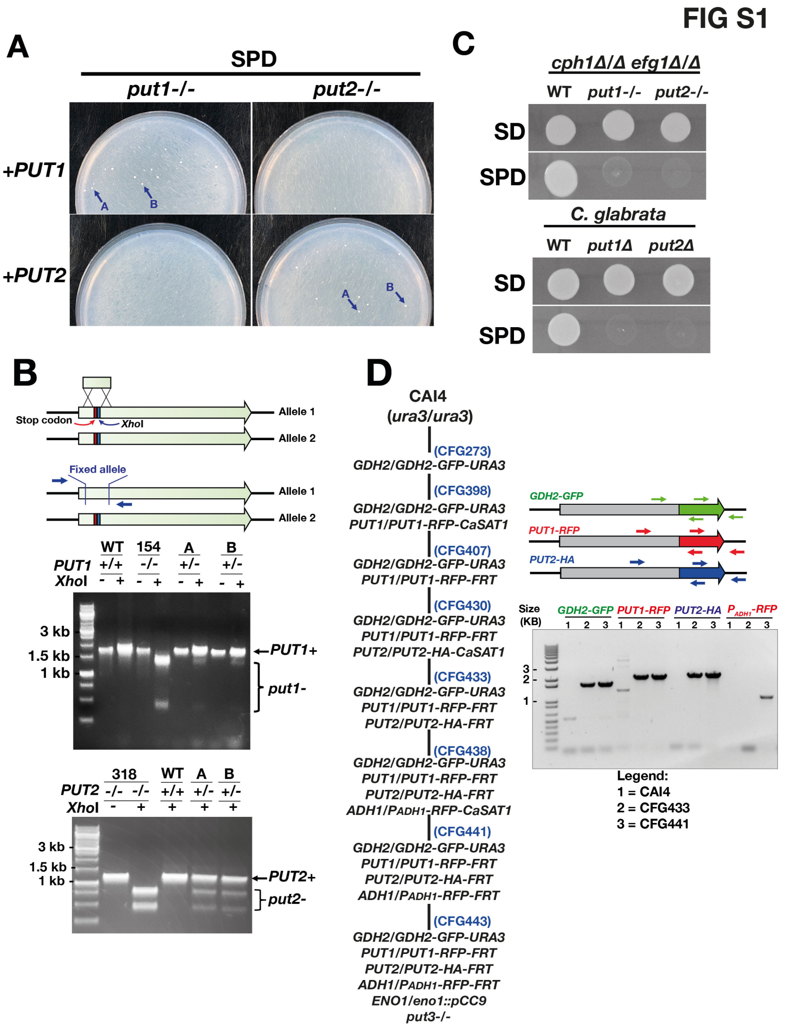


**Figure S1. Strain construction and targeted reconstitution**

(**A**) Targeted reconstitution: *put1-/-* (CFG154) and *put2-/-* (CFG318) strains were transformed with wildtype *PUT1* and *PUT2* fragments, respectively, and proline-utilizing (Put$^{+}$) colonies were selected on SPD. As control, the *PUT2* and *PUT1* fragments were introduced into the *put1-/-* and *put2-/-* strains, respectively; no transformants were obtained. (**B**) Verification of the reconstructed *PUT1* and *PUT2* alleles. The colonies with arrows (**A**) were purified and their genomes analyzed by PCR-RD. The heterozygosity at the indicated gene locus was confirmed, i.e., *PUT1+/- (PUT1+/put1-)* and *PUT2*+/- (*PUT2+/put2-*). Primers (shown in blue) facilitate the amplification of both the wildtype Alleles 1 and *Xho*I containing CRISPR/Cas9 inactivated Alleles 2. The amplified fragments were digested with *Xho*I and fragment lengths were analyzed by electrophoresis (1% agarose gel). The fragments with the inactivated alleles are cleaved by *Xho*I resulting in two bands (indicated by the brackets), whereas the reconstructed wildtype fragment is refractory to *Xho*I digestion and runs as a single band (arrow). (**C**) *C. albicans cph1*Δ/Δ *efg1*Δ/Δ (WT, CASJ041), *cph1*Δ/Δ *efg1*Δ/Δ *put1-/-* (CFG344), *cph1*Δ/Δ *efg1*Δ/Δ *put2-/-* (CFG345) and *C. glabrata* (WT, CBS138), Cg*put1*Δ (GFS003), Cg*put2*Δ (GFS005) strains were spotted on non-selective (SD) and selective (SPD) media as indicated. The inactivation of *PUT* genes resulted in the lack of growth on selective media, indicating that inability of the mutants to use catabolize proline. (**D**) Schematic diagram of steps required to construct the triply-tagged reporter strain. The starting parental strain was CAI4 (*ura3/ura3*); the complete genotypes of the strains are listed in Methods. PCR-verification of reporter constructs used two pairs of primer pairs described in Methods. Sizes of fragments amplified with primer pairs flanking the tag inserts: *GDH2-GFP* (1.6 kb), *PUT1-RFP* (2.1 kb), *PUT2-HA* (2.3 kb), $P_{ADH1}$-*RFP* (1.2 kb).

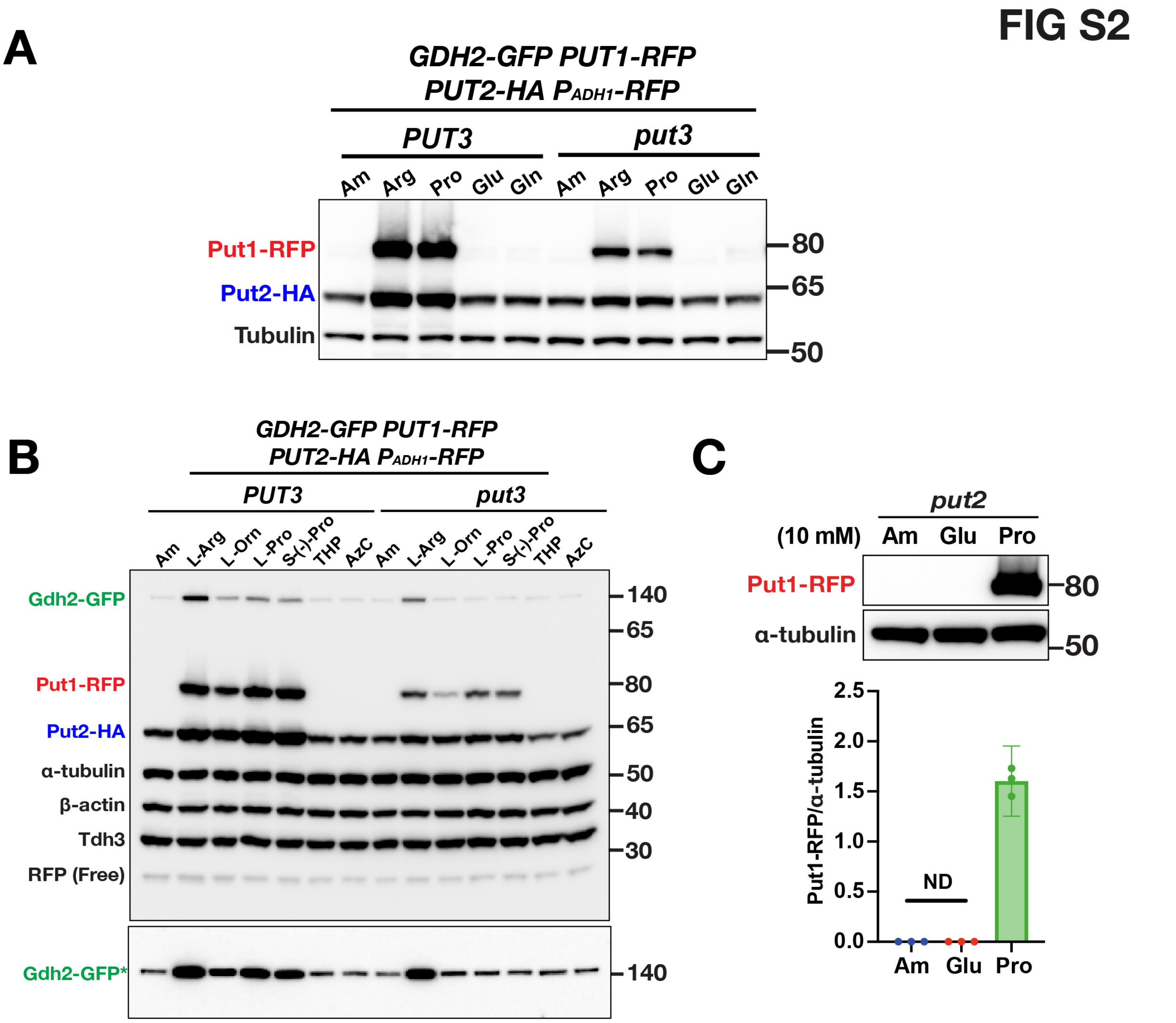


**Figure S2. Put3-dependent and -independent induction of PUT enzymes**
(**A**) Exponentially growing cultures of triple tagged reporter strains CFG441 (*PUT3*) and CFG443 (*put3*) in SGL were induced with 10 mM of the indicated nitrogen source for 1 h. Cell extracts were prepared and the expression of the Put1, Put2 and tubulin was analyzed by immunoblot using an optimized antibody cocktail (see Methods). Note that Put1-RFP is not detected in cultures containing either glutamate or glutamine. (**B**) Specificity of Put3 to proline. Strains, growth and analysis as in (A) of cultures induced with the addition of 10 mM of the indicated compounds: Ammonium sulfate (Am); L-ornithine; L-arginine; L-proline; S-(-)-proline; T-4-hydroxy-L-proline; and Azetidine carboxylate (AzC). Gdh2-GFP signals (*) were separately enhanced (lower panel) via the high slider in Image Lab (BioRad). (**C**) Glutamate is not metabolized to proline. Strain CFG469 was grown and induced as in (A) with 10 mM ammonium sulfate (Am), glutamate or proline as indicated. Note that Put1-RFP was not detected (ND) in Am or glutamate induced cultures. The immunoblots are representative of at least three independent experiments.

FIG S3

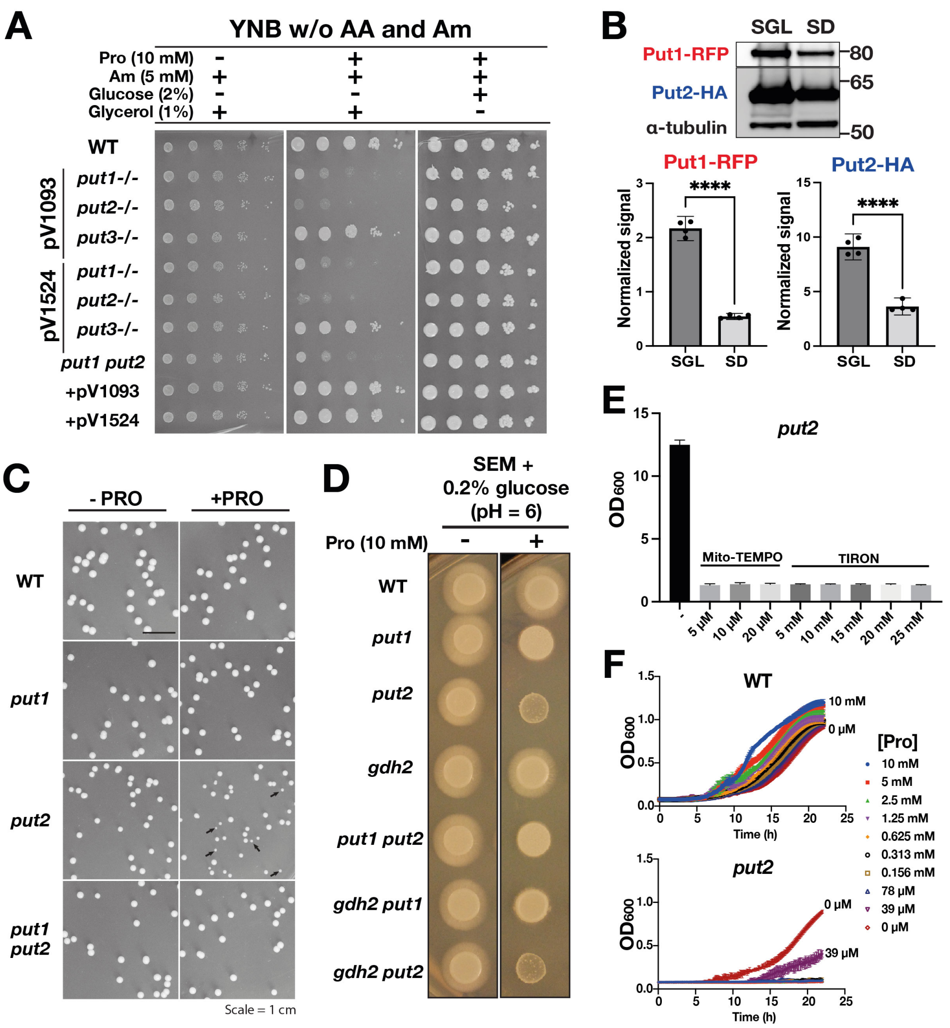


**Figure S3. Proline catabolic mutants are sensitive to exogenous proline**

(**A**) Serially diluted *C. albicans* cells of the indicated genotypes (same as in Fig. 1C**)** were spotted onto buffered synthetic minimal medium (pH = 6) containing 5 mM ammonium sulfate (Am) as nitrogen source and either 1% glycerol or 2% glucose as carbon source. Excess proline (10 mM) was added as indicated. Plates were photographed after 4 days of growth at 30 °C. (**B**) Strain CFG433, grown to log phase in SGL or SD, was induced with 10 mM proline for 1 h, and cell extracts were analyzed by immunoblot. The signals from Put1-RFP and Put2-HA were normalized to α-tubulin. Data presented are from 4 biological replicates (Ave. with 95% CI; *****p*<0.0001 by student *t*-test). (**C**) Cells from 72 h-old SD cultures of the indicated strains (same as in Fig. 3A**)** were plated for single colonies on YPD and grown for 2 days. The colonies from the *put2* mutant are heterogenous in size, both large and small colonies (black arrows) are evident. Images were representative of at least 3 biological replicates. Scale 1 = cm. (**D**) Five µl of cell suspensions of the indicated strains (same as in Fig. 3A**)** were spotted on buffered SEM medium (pH =6) containing 10 mM glutamate with and without 10 mM proline as indicated. The resulting macro-colonies were photographed after 72 h at 30 °C. (**E**) ROS (superoxide) scavengers failed to rescue the growth of *put2* in the presence of exogenous proline. CFG318 was grown as in Fig. 3D with and without the indicated concentrations of Mito-TEMPO or TIRON. The results are the average of at 4 biological replicates (with 95% CI). (**F**) Growth of both WT (SC5314) and *put2* mutant (CFG318) strains grown in SGL medium in the presence of the indicated concentration of proline. Cells were grown in a 96-well TECAN microplate as in Fig. 3F.

**FIG S4**

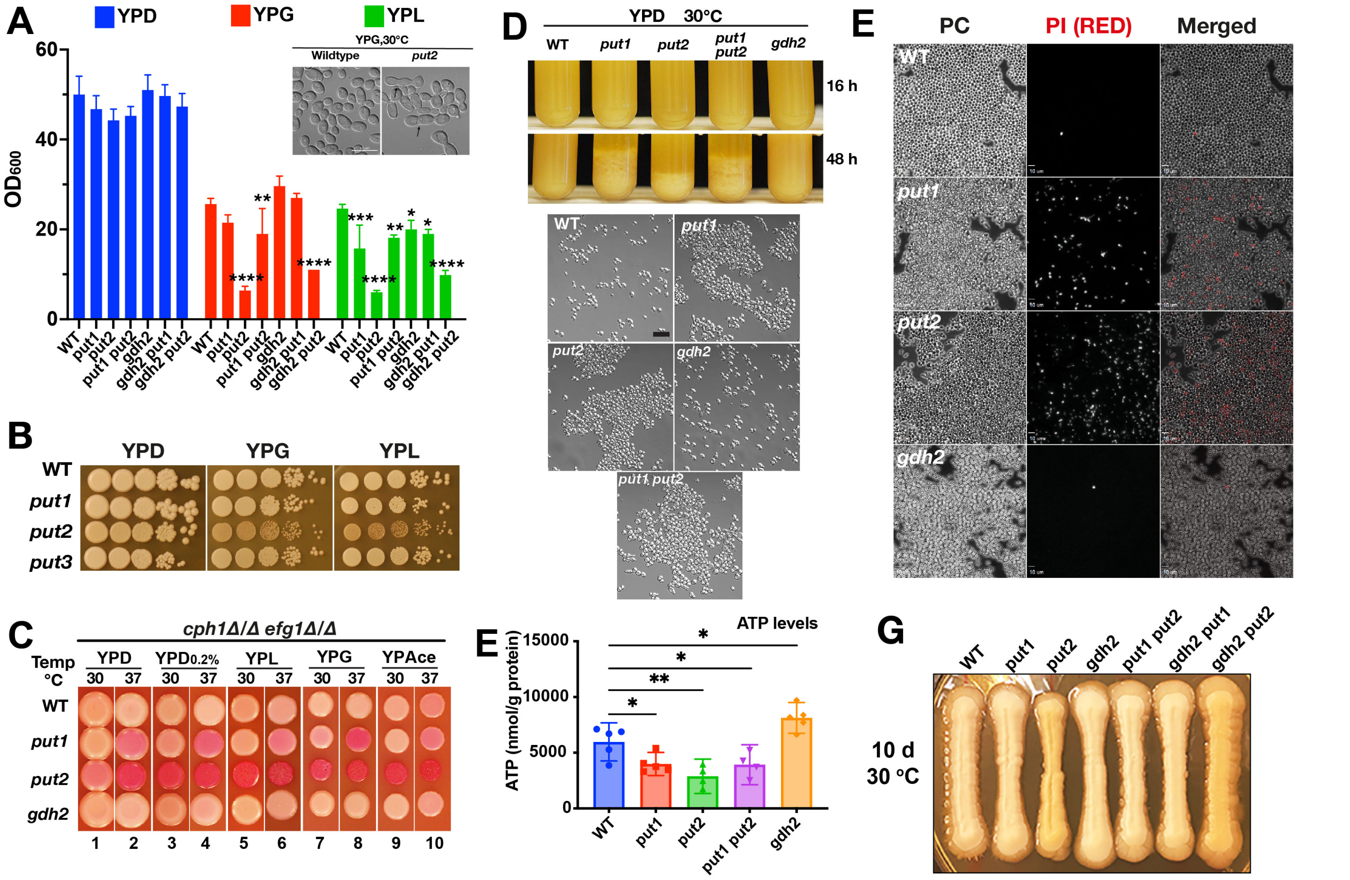


**Figure S4. Cells lacking the capacity to catabolize proline exhibit enhanced death.**
(**A**) Growth of *put* mutants in media with different carbon sources. Cells collected from YPD pre-cultures were washed and inoculated into tubes with YP containing the indicated carbon source at an $OD_{600} \approx 0.01$, growth was assessed after 24 h at 30 °C. Note that *put2* cells grown in YPG have noticeably higher number of cells forming trimera a phenotype indicative of stress (black arrows, inset). Results, presented per carbon source, were derived from 4 biological replicates (Ave.±SD; one-way ANOVA (YPD, $p = 0.0398$; YPG, YPL, and YPAce, $p < 0.0001$) with Dunnett's post hoc test relative to wildtype (****$p$ <0.0001, ***$p$ <0.001, **$p$ <0.01, *$p$ <0.05). Strains used: WT (SC5314); *put1* (CFG154), *put2* (CFG318); *put3* (CFG156); *put1 put2* (CFG159); *gdh2* (CFG279); *gdh2 put1* (CFG364); *gdh2 put2* (CFG366). (**B**) Ten-fold serial dilutions of cell suspensions ($OD_{600} \approx 1$) prepared from cultures as in (A) were spotted on YPD, YPG or YPL. The plates were incubated 2 days at 30 °C and photographed. (**C**) Five microliters of *cph1*Δ/Δ *efg1*Δ/Δ (CASJ041) or its derivatives (*put1*, CFG344; *put2*, CFG345; *gdh2*, CFG352) ($OD_{600} \approx 1.0$) were spotted on the indicated plates containing Phloxine B (10 µg/ml), and incubated for 3 days at 30 or 37 °C as indicted and photographed. (**D**) Cells were grown in liquid YPD and flocculation was assessed after 16 and 48 h of growth at 30 ºC as indicated. Cultures were vigorously vortexed and let stand immobile for 3 min and photographed. (**E**) Intracellular ATP of *put* mutants entering the saturated phase is lower than wildtype. ATP was extracted from cells grown 16 h in liquid YPD as in (B) and harvested. ATP concentrations, derived from at least 3 biological replicates, are presented normalized to total protein content (Error bars, 95% CI; *$p$ <0.05, **$p$ <0.01 by one-way ANOVA with Dunnett's post hoc test). (**F**) Propidium iodide (PI) staining of cells from 48 h-old cultures of wildtype (SC5314), *put1* (CFG154), *put2* (CFG318), *put3* (CFG156), *put1 put2* (CFG159), and *gdh2* (CFG279). Results are representative of at least three independent experiments. (**G**) Cells of the indicated genotypes were streaked on YPD, the plate was incubated 10 days at 30 °C and photographed. Note the increase in yellow hue in *put2* and *gdh2 put2* strains.

FIG S5

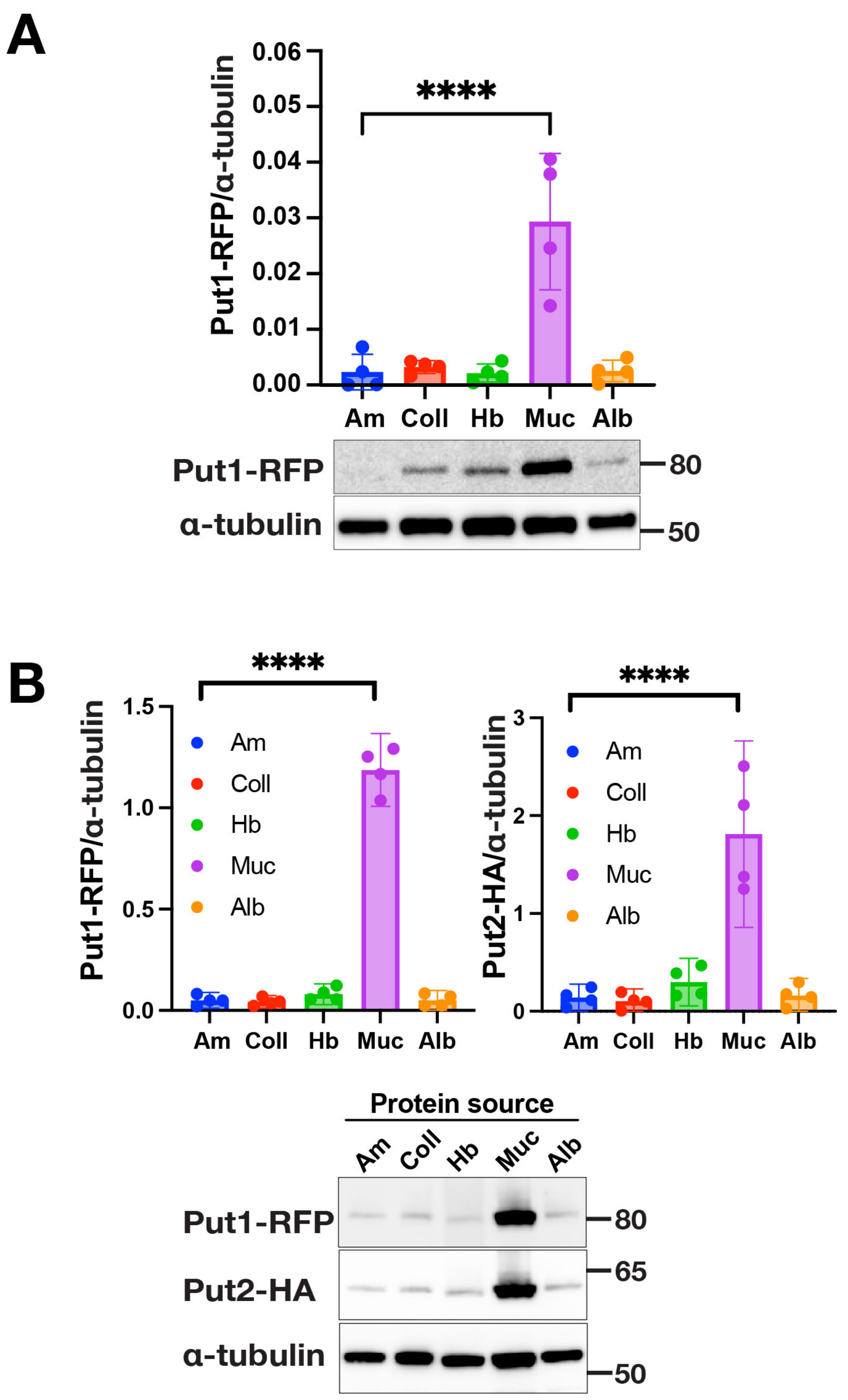


**Figure S5. Expression of Proline Utilization enzymes during growth in the presence of different protein sources**
CFG433 cells were pre-grown in SGL, washed, and used to inoculate modified HBSS medium containing glucose and lactate plus the indicated protein as sole nitrogen source (0.5 mg/ml) and incubated at 37 °C for 24 h (**A**) and 72 h (**B**). Cell extracts were prepared and analyzed by immunoblot and developed to detect Put1 and Put2 as indicated. Legend: Am (Ammonium sulfate), Coll (Collagen), Hb (Hemoglobin), Muc (Mucin), Alb (Human serum albumin). Results (Ave.±SD, n = 4) were analyzed by one-way ANOVA with Dunnett's post hoc test relative to Am (*****p* <0.0001). Proline derived from mucin appears to be easily assimilated by *C. albicans*.

### FIG S6

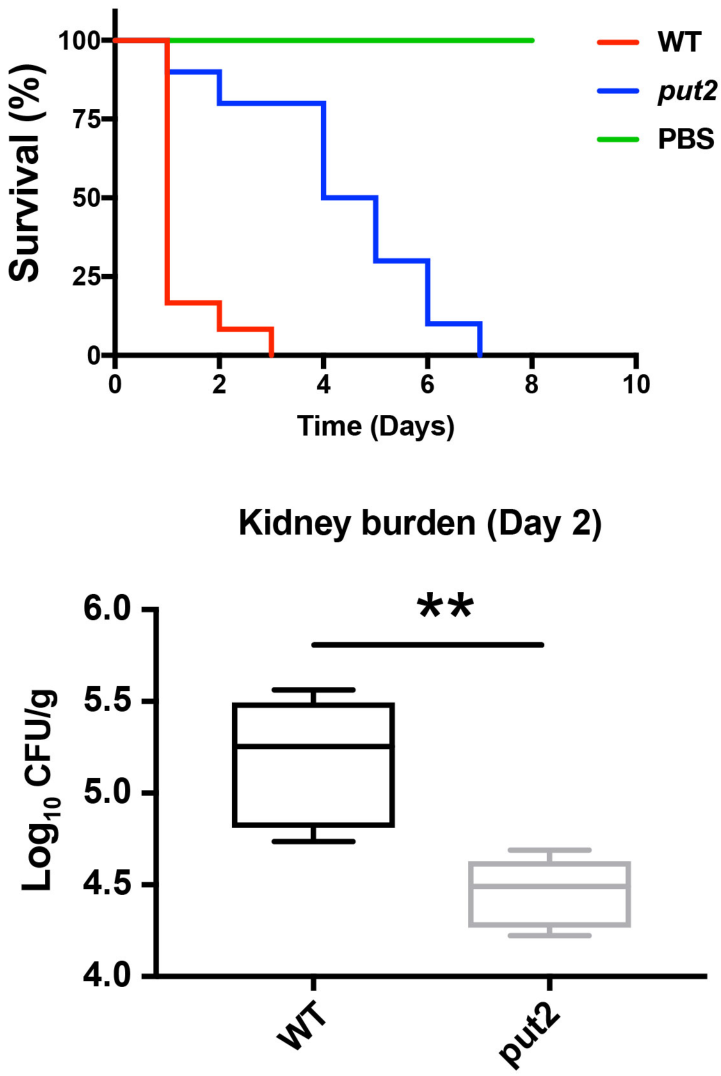


**Figure S6. Mutations inactivating proline catabolism attenuate virulence in BALB/cAnNCrl mice**
Upper panel, female BALB/cAnNCrl mice were infected via the lateral tail vain with 5x10$^5$ CFU of *C. albicans* wildtype (SC5314) or *put2* (CFG318) mutant. Each curve in the plot are the average of 3 independent experiments (10 mice/strain). Mice infected with *put2-/-* survived longer compared to wildtype (*****p*<0.0001 by Log-rank (Mantel-Cox) test). Lower panel, the fungal burden in kidneys extracted from mice 2 days after infection. Box and whiskers plot showing significantly lower fungal burden in the kidney of mice infected with *put2* mutant compared to wildtype (***p*=0.0039 by student *t*-test).

| **Table S1. Key reagents and resources** | | |
|---|---|---|
| **Reagent or Resource** | **Source** | **Identifier** |
| **Antibodies** | | |
| Anti-ATP5A [EPR13030(B)], rabbit monoclonal | Abcam | Cat#ab176569 |
| Anti-β-actin, mouse monoclonal | Abcam | Cat#ab8224 |
| Anti-GFP, Living Colors® A.v. (JL-8), mouse monoclonal | Takara | Cat#632381 |
| Anti-HA-Peroxidase, High Affinity (clone 3F10) rat monoclonal | Roche | Cat#12013819001 |
| Anti-mCherry, rabbit polyclonal | Abcam | Cat#ab167453 |
| Anti-GAPDH [GT239], mouse monoclonal | Genetex | Cat#GTX627408 |
| Anti-tubulin [YOL1/34] conjugated to HRP, rat monoclonal | Abcam | Cat#ab196583 |
| Goat anti-mouse IgG (H+L) secondary antibody, poly-HRP | Invitrogen | Cat#31430 |
| Goat anti-rabbit IgG (H+L) secondary antibody, poly-HRP | Invitrogen | Cat#31460 |
| **Chemicals, peptides, and recombinant proteins** | | |
| 2-aminobenzaldehyde (O-ABZ) | Sigma-Aldrich | Cat#A9628 |
| Agar | Formedium | Cat#A9628 |
| Albumin from human serum | Sigma-Aldrich | Cat#A1653 |
| Ammonium sulfate | VWR | Cat#21333.296 |
| Carbenicillin disodium salt | AppliChem GmbH | Cat#A1491 |
| cis-4-Hydroxy-L-proline | Sigma-Aldrich | Cat#H1637 |
| cis-4-Hydroxy-D-proline | Sigma-Aldrich | Cat#H5877 |
| Collagen from bovine achilles tendon | Sigma-Aldrich | Cat#C9879 |
| Collagen from human placenta Bornstein and Traub Type IV | Sigma-Aldrich | Cat#C7521 |
| cOmplete Mini, EDTA free (protease inhibitor cocktail) | Roche | Cat#11836170001 |
| D-(+)-Glucose | Sigma-Aldrich | Cat#G7528 |
| D-Proline | Sigma-Aldrich | Cat#858919 |
| DL-Lactic acid, ~90% (T) | Sigma-Aldrich | Cat#69785 |
| DTT - DL-1,4-Dithiothreitol | Acros Organics | Cat#165680050 |
| DreamTaq Green DNA Polymerase | Thermo Fisher Scientific | Cat#EP0702 |
| ExTaq ® Hot Start Version (250U) | Takara | Cat#RR006A |
| Fastdigest *Kpn*I | Thermo Fisher Scientific | Cat#FD0524 |
| Fastdigest *Nco*I | Thermo Fisher Scientific | Cat#FD0573 |
| Fastdigest *Pvu*I | Thermo Fisher Scientific | Cat#FD0624 |

| | | |
|---|---|---|
| Fastdigest *Sac*I | Thermo Fisher Scientific | Cat#FD1133 |
| Fastdigest *Xho*I | Thermo Fisher Scientific | Cat#FD0694 |
| FITC - Fluorescein-5-isothiocyanate | Sigma-Aldrich | Cat#F7250 |
| Gibson Assembly® Master Mix | New England Biolabs | Cat# E2611 |
| Glycerol, ≥99.0% (GC) | Sigma-Aldrich | Cat#G7757 |
| Hemoglobin from bovine blood | Fluka | Cat#8449 |
| Histopaque-1119 | Sigma-Aldrich | Cat# 11191 |
| Isoflurane, Attane vet 1000 mg/g | Piramal Healthcare UK Ltd. | Cat#NDC66794017-25 |
| L-Arginine monohydrochloride | Sigma-Aldrich | Cat#A5131 |
| L-Azetidine-2-Carboxylic acid | Sigma-Aldrich | Cat#A0760 |
| L-Glutamic acid | Sigma-Aldrich | Cat# G1251 |
| L-Glutamine | Sigma-Aldrich | Cat#G3126 |
| L-Ornithine | Sigma-Aldrich | Cat#O2375 |
| L-Proline | Sigma-Aldrich | Cat#P0380 |
| Luminol | Sigma-Aldrich | Cat#123072 |
| MitoSOX™ Red | Invitrogen | Cat#M36008 |
| MitoTEMPO | Sigma-Aldrich | Cat#SML0737 |
| Mucin from porcine stomach Type II | Sigma-Aldrich | Cat#M2378 |
| N-acetyl-cysteine (NAC) | Sigma-Aldrich | Cat#A7250 |
| Nourseothricin (clonNAT) | Jena Bioscience | Cat#AB-102XL |
| NuPAGE™ 3 to 8%, Tris-Acetate, 1.0 mm, Mini Protein Gel | Thermo Fisher Scientific | Cat#EA03752BOX |
| Peptone | Formedium | Cat#PEP03 |
| Peptone | OXOID | Cat#PEP03 |
| Peroxidase from horseradish | Sigma-Aldrich | Cat#P8125-5K |
| Phloxine B, 85%, high purity biological stain | VWR | Cat#ACRO189470050 |
| Phusion™ High-Fidelity DNA Polymerase | Thermo Fisher Scientific | Cat#F530L |
| Propidium iodide (1.0 mg/ml in $H_2O$) | Thermo Fisher Scientific | Cat#P3566 |
| PureCol™ EZ gel | Sigma-Aldrich | Cat#5074 |
| (S)-(-)-Proline | Sigma-Aldrich | Cat#8160190025 |
| Sabouraud Dextrose Agar Medium | ACMEC biochemical | Cat#AC15825 |
| SDS - Sodium dodecyl sulfate | Sigma-Aldrich | Cat#75746 |
| SuperSignal Dura West Extended Duration Substrate | Thermo Fisher Scientific | Cat#34076 |
| Thiazolidine-2-carboxylic acid | Sigma-Aldrich | Cat#467995 |

| | | |
|---|---|---|
| TIRON - 4,5-Dihydroxy-1,3-benzenedisulfonic acid disodium salt monohydrate | Sigma-Aldrich | Cat#172553 |
| trans-4-Hydroxy-L-proline | Sigma-Aldrich | Cat#H54409 |
| Yeast extract powder | Formedium | Cat#YEA02 |
| Yeast extract powder | OXOID | Cat#LP0021 |
| Yeast Nitrogen Base without amino acids and ammonium sulfate | BD Difco | Cat#11743014 |
| **Critical commercial assays** | | |
| Molecular Probes™ ATP Determination Kit | Thermo Fisher Scientific | Cat#A22066 |
| GeneJet PCR Purification Kit | Thermo Fisher Scientific | Cat#K0702 |
| GeneJet Plasmid Miniprep Kit | Thermo Fisher Scientific | Cat#K0503 |
| Masterpure™ Yeast DNA Purification Kit | Epicentre | Cat#MPY80200 |
| QIAquick Gel Extraction Kit | Qiagen | Cat#28706 |
| **Recombinant DNA** | | |
| Plasmid: pFA-GFPγ-URA3 | (*18*) | pFA-GFPγ-URA3 |
| Plasmid: pFA6a-3xHA-SAT1-FLP | (*19*) | pFA6a-3xHA-SAT1-FLP |
| Plasmid: pJA21 | (*1*) | $P_{ADH1}$-*RFP-CaSAT1* flipper cassette |
| Plasmid: pV1093 | (*2*) | CRISPR/Cas9 cassette, general vector |
| Plasmid: pV1524 | (*3*) | CRISPR/Cas9 cassette, general vector |
| Plasmid: pFS080 | (*1*) | *PUT1* sgRNA inserted into pV1093 |
| Plasmid: pFS083 | (*1*) | *PUT2* sgRNA inserted into pV1093 |
| Plasmid: pFS084 | (*1*) | *PUT3* sgRNA inserted into pV1093 |
| Plasmid: pFS088 | (*1*) | *PUT1* sgRNA inserted into pV1524 |
| Plasmid: pFS092 | (*1*) | *PUT2* sgRNA inserted into pV1524 |
| Plasmid: pFS090 | (*1*) | *PUT3* sgRNA inserted into pV1524 |
| Plasmid: pSFS3b | (*7*) | Donor plasmid for SAT1 flipper fragment used for Gibson assembly |
| Plasmid: YEp352 | (*20*) | Donor plasmid for *E. coli* Ori and $Amp^R$ fragment for Gibson assembly |
| Plasmid: YEp352-*NAT1*-Cg*PUT1*urdr | This study | Plasmid for *C. glabrata PUT1* gene deletion |
| Plasmid: YEp352-*NAT1*-Cg*PUT2*urdr | This study | Plasmid for *C. glabrata PUT2* gene deletion |
| **Experimental models: Organisms/Strains** | | |
| **Mouse strains** | | |
| C57BL/6 | Zhejiang Vital River Laboratory Animal Technology Co., Ltd. | |
| BALB/cAnNCrl | Charles River, Germany | |

| | | |
|---|---|---|
| BALB/cByJ SOPF | Charles River, France | |
| ***Drosophila melanogaster* strains** | | |
| *Bom*$^{\Delta 55C}$ | (*21*) | Toll-regulated Bomanin effectors-deficient flies |
| ***C. albicans* CAI4-derived strains** | | |
| *C. albicans:* PLC016, PMRCA18 | (*22*) | *ura3::imm434/ura3::URA3* |
| *C. albicans:* CFG219 | (*8*) | *ura3::imm434/ura3::imm434 iro1/iro1::imm434 PUT2/PUT2-GFP-URA3* |
| *C. albicans:* CFG237 | (*8*) | *ura3::imm434/ura3::imm434 iro1/iro1::imm434 PUT2/PUT2-GFP-URA3 ADH1/adh1::P$_{ADH1}$-RFP-CaSAT1* |
| *C. albicans:* CFG259 | (*8*) | *ura3::imm434/ura3::imm434 iro1/iro1::imm434 PUT2/PUT2-GFP-URA3 ADH1/adh1::P$_{ADH1}$-RFP-FRT* |
| *C. albicans:* CFG301 | This work | *ura3::imm434/ura3::imm434 iro1/iro1::imm434 PUT2/PUT2-GFP-URA3 ADH1/adh1::P$_{ADH1}$-RFP-FRT ENO1/eno1::P$_{ENO1}$-CC9-pFS084 put3-/-* |
| *C. albicans:* CFG407 | (*8*) | *ura3::imm434/ura3::imm434 iro1/iro1::imm434 GDH2/GDH2-GFP-URA3 PUT1/PUT1-RFP-FRT-FRT* |
| *C. albicans:* CFG430 | This work | *ura3::imm434/ura3::imm434 iro1/iro1::imm434 GDH2/GDH2-GFP-URA3 PUT1/PUT1-RFP-FRT-FRT PUT2/PUT2-HA-CaSAT1* |
| *C. albicans:* CFG433 | This work | *ura3::imm434/ura3::imm434 iro1/iro1::imm434 GDH2/GDH2-GFP-URA3 PUT1/PUT1-RFP-FRT-FRT PUT2/PUT2-HA-FRT* |
| *C. albicans:* CFG438 | This work | *ura3::imm434/ura3::imm434 iro1/iro1::imm434 GDH2/GDH2-GFP-URA3 PUT1/PUT1-RFP-FRT-FRT PUT2/PUT2-HA-FRT ADH1/adh1::P$_{ADH1}$-RFP-CaSAT1* |
| *C. albicans:* CFG441 | This work | *ura3::imm434/ura3::imm434 iro1/iro1::imm434 GDH2/GDH2-GFP-URA3 PUT1/PUT1-RFP-FRT-FRT PUT2/PUT2-HA-FRT ADH1/adh1::P$_{ADH1}$-RFP-FRT* |
| *C. albicans:* CFG443 | This work | *ura3::imm434/ura3::imm434 iro1/iro1::imm434 GDH2/GDH2-GFP-URA3 PUT1/PUT1-RFP-FRT-FRT PUT2/PUT2-HA-FRT ADH1/adh1::P$_{ADH1}$-RFP-FRT ENO1/eno1::P$_{ENO1}$-CC9-pFS084 put3-/-* |
| ***C. albicans* SC5314-derived strains** | | |
| *C. albicans*: SC5314, PLC005 | (*23*) | Prototrophic wildtype |
| *C. albicans*: CASJ041 | (*14*); KK Collection | *cph1Δ::FRT/ cph1Δ::FRT efg1Δ::FRT/efg1Δ::FRT* |
| *C. albicans*: CFG143 | (*1*) | *ENO1/eno1::P$_{ENO1}$-CC9-pFS083 put2-/-* |
| *C. albicans*: CFG146 | (*1*) | *NEUT5/neut5::P$_{ENO1}$-CC9-pFS090 put3- /-* |
| *C. albicans*: CFG149 | (*1*) | *ENO1/eno1::P$_{ENO1}$-CC9-pFS080 put1-/-* |
| *C. albicans*: CFG150 | (*1*) | *ENO1/eno1::P$_{ENO1}$-CC9-pFS084 put3-/-* |

| | | |
|---|---|---|
| *C. albicans*: CFG154 | (*1*) | *NEUT5/neut5::FRT put1-/-* |
| *C. albicans:* CFG156 | (*1*) | *NEUT5/neut5::FRT put3-/-* |
| *C. albicans*: CFG159 | (*1*) | *NEUT5/neut5::FRT put1-/-* *ENO1/eno1::*$P_{ENO1}$*-CC9-pFS083 put2-/-* |
| *C. albicans*: CFG181 | (*1*) | *ENO1/eno1::*$P_{ENO1}$*-CC9-pV1093* |
| *C. albicans*: CFG182 | (*1*) | *NEUT5/neut5::*$P_{ENO1}$*-CC9-pV1524* |
| *C. albicans*: CFG187 | This work | *PUT3/PUT3-HA-CaSAT1 (Clone 1)* |
| *C. albicans*: CFG188 | This work | *PUT3/PUT3-HA-CaSAT1 (Clone 2)* |
| *C. albicans*: CFG318 | (*1*) | *NEUT5/neut5::FRT put2-/-* |
| *C. albicans*: CFG344 | This work | *cph1Δ::FRT/cph1Δ::FRT efg1Δ::FRT/efg1Δ::FRT NEUT5/neut5::FRT put1-/-* |
| *C. albicans*: CFG345 | This work | *cph1Δ::FRT/cph1Δ::FRT efg1Δ::FRT/efg1Δ::FRT NEUT5/neut5::FRT put2-/-* |
| *C. albicans*: CFG352 | (*8*) | *cph1Δ::FRT/cph1Δ::FRT efg1Δ::FRT/efg1Δ::FRT NEUT5/neut5::FRT gdh2-/-* |
| *C. albicans*: CFG364 | This work | *NEUT5/neut5::FRT gdh2-/-* *ENO1/eno1::*$P_{ENO1}$*-CC9-pFS080 put1-/-* |
| *C. albicans*: CFG366 | This work | *NEUT5/neut5::FRT gdh2-/-* *ENO1/eno1::*$P_{ENO1}$*-CC9-pFS083 put2-/-* |
| *C. albicans*: CFG379 | This work | *NEUT5/neut5::FRT PUT1+/ put1-* |
| *C. albicans*: CFG380 | This work | *NEUT5/neut5::FRT PUT1+/ put1-* |
| *C. albicans*: CFG381 | This work | *NEUT5/neut5::FRT PUT2+/ put2-* |
| *C. albicans*: CFG382 | This work | *NEUT5/neut5::FRT PUT2+/ put2-* |
| ***C. albicans* BWP17-derived strains** | | |
| *C. albicans*: PLC096 | (*24*) | *ura3::imm434/ura3::imm434 iro1/iro1::imm434 his1::hisG/his1::hisG arg4/arg4 ADH1/*$P_{ADH1}$*-yEmRFP-URA3 NEUT5/neut5::*$P_{ENO1}$*-CC9-pFS092 put2-/-* |
| *C. albicans*: CFG474 | This work | *ura3::imm434/ura3::imm434 iro1/iro1::imm434 his1::hisG/his1::hisG arg4/arg4 ADH1/*$P_{ADH1}$*-yEmRFP-URA3 NEUT5/neut5::*$P_{ENO1}$*-CC9-pFS092 put2-/-* |
| *C. albicans*: CFG479 | This work | *ura3::imm434/ura3::imm434 iro1/iro1::imm434 his1::hisG/his1::hisG arg4/arg4 ADH1/*$P_{ADH1}$*-yEmRFP-URA3 NEUT5/neut5::FRT put2-/-* |
| ***C. glabrata strains*** | | |
| *C. glabrata*: ATCC2001/CBS138 | ATCC | Prototrophic wildtype |
| *C. glabrata*: GFS003 | This work | *put1Δ::FRT* |
| *C. glabrata*: GFS005 | This work | *put2Δ::FRT* |
| **Other yeast strains** | | |
| *C. albicans*: MAY7 | (*25*); MA Collection | Clinical isolate; fluconazole resistant |
| *C. albicans*: PLC124 | POL Collection | From Karolinska Hospital (Solna) isolated from male patient with knee prosthesis |
| *C. utilis*: F608 | POL Collection | Prototrophic wildtype |

| | | |
|---|---|---|
| *C. glabrata*: Peu927 | OB Collection | Clinical isolate |
| *C. tropicalis*: SM1541 | UR Collection | Clinical isolate |
| *C. tropicalis*: ATCC750 (1) | VDV Collection | Clinical isolate |
| *C. tropicalis*: ATCC750 (2) | UR Collection | Clinical isolate |
| *C. dubliniensis:* SMI718 | UR Collection | Clinical isolate |
| *C. dubliniensis*: Wü284 (1) | (*26*); SR Collection | Clinical isolate |
| *C. dubliniensis*: Wü284 (2) | (*26*); CU Collection | Clinical isolate |
| *C. parapsilosis*: ATCC22019 (1) | UR Collection | Clinical isolate |
| *C. parapsilosis*: ATCC22019 (2) | SR Collection | Clinical isolate |
| *C. parapsilosis*: ATCC22019 (3) | VDV Collection | Clinical isolate |
| *C. krusei*: ATCC6258 | SR Collection | Clinical isolate |
| *C. auris*: CBS10913 | (*27*); KK Collection | Clinical isolate |
| *C. lusitaniae*: DSM 70102 | KK Collection | Clinical isolate |
| *C. guilliermondii*: ATCC6260 | KK Collection | Clinical isolate |
| *S. cerevisiae*: S288c | Reference strain | Prototrophic haploid (1N) |
| *S. cerevisiae*: KRY001 | (*16*) | Σ1278b-derived diploid (2N) |
| *S. cerevisiae*: CFG638 | POL Collection | Clinical isolate from Karolinska Hospital (Huddinge), Sweden |
| *S. cerevisiae*: CFG639 | POL Collection | Clinical isolate from Karolinska Hospital (Huddinge), Sweden |
| *S. cerevisiae*: CFG640 | POL Collection | Clinical isolate from Karolinska Hospital (Huddinge), Sweden |
| *S. cerevisiae*: CFG641 | POL Collection | Clinical isolate from Karolinska Hospital (Huddinge), Sweden |
| *Cryptococcus neoformans* | KK Collection | Clinical isolate |
| **Oligonucleotides** | | |
| Oligonucleotides are described in **Table S2** | N/A | N/A |
| **Software and algorithms** | | |
| GraphPad Prism version 9 | GraphPad | GraphPad Software; https://www.graphpad.com; RRID: SCR_002798 |
| Huygens Deconvolution | Scientific Volume Imaging | https://svi.nl/HomePage |
| ImageJ software (Fiji v. 2.0.0) | NIH | https://imagej.net/ |
| Image Lab 6.1.0 | Bio-Rad | https://www.bio-rad.com |
| LAS-X | Leica microsystems | |
| OxyTrace+ Windows® software | Hansatech Instruments | |
| Serial Cloner 2.6 | Serial Cloner | http://serialbasics.free.fr/Serial_Cloner.html |
| SnapGene 6.1.2 | Dotmatics | https://www.snapgene.com |
| Zen Blue software | Zeiss | |

**Table S2. Oligonucleotides**

| No | Primer Name | Sequence | Reference |
|---|---|---|---|

| | | | |
|---|---|---|---|
| 1 | 161-PUT1-VerF | CATCGGTTATTATTCTTCTTG | (*1*) |
| 2 | 162-PUT1-VerR | GTTTAACCACTTCCAAATAATC | (*1*) |
| 3 | 244-PUT2-VerF | CTAGCGGATTAACTATTCGC | (*1*) |
| 4 | 245-PUT2-VerR | GGATAATGCGGCTGTAGCAG | (*1*) |
| 5 | 237-PUT3-VerF | CGTGCATTACTTCATGTAATC | (*1*) |
| 6 | 238-PUT3-VerR | GGACGAAGGTATTGTTTGAGG | (*1*) |
| 7 | 377-PUT1ver2 | CGAGACAATCAATGGAAACC | This work |
| 8 | 378-PUT2ver2 | CCACAGACTGAATGAAGAAG | This work |
| 9 | FS95 | GGCATAGCTGAAACTTCGGC | (*1*) |
| 10 | 275-C-HATagPUT2F | CTGGTAGTGGTAACATTTTATCCAGATTTGTTTCTATTAGAAACATTAAAGAAAACTTTTACGAATTGACTGATTTCAAATATCCATCCAATTATCAAAACATCTTTTACCCATACGATG | (*1*) |
| 11 | 276-C-HATagPUT2R | GGAAACAACATGAACACCTTATGTAAGAAAACCTCTTCTTAATATAAATATTTACATTCACACATTAACTATATAAAGTAATAACTAATCTCGTTTCTCGCAGGTTAACCTGGCTTATCG | (*1*) |
| 12 | P110_FS244HAFOR | CTAGCGGATTAACTATTCGC | (*1*) |
| 13 | P111_FS340HAREV | CGTCATATGGATAGGATCCTG | (*1*) |
| 14 | 231PUT1Upstr | CCACTCAATCAATGATCATCC | This work |
| 15 | FS321ADH1upstr | GAGACCCAATGCAAAGCCAG | This work |
| 16 | JAV35-3'RFPtest | GAATCTTGAGTAACAGTAAC | This work |
| 17 | 339VerGDH2HAF | GTTCCATGTGGTGGTAGACC | (*8*) |
| 18 | 379GFPuniv | GTCCATCTTTACAGTCCTGTC | (*8*) |
| 19 | 277C-HATagPUT3F | TTATATTTAATACTGCATCGCCTGATGTTGGCACTAGTGTCCATTCAGGGTATTCAAACTTGATAAACCATGAATTTCAAGATTTCATGGATCAATCTAACATCTTTTACCCATACGATG | This work |
| 20 | 278C-HATagPUT3R | GTTTTTGTAATATATTGTATTATATAGAAAATTTTATTACCATCACAGAATAAATGTACAGACATAAATATATATTTGCCTCACTCCCGCACAATCACTGCAGGTTAACCTGGCTTATCG | This work |
| 21 | RT-PUT1Top | GATTATATTCTAAACAATCTTTAAATACATTCAAAAAAGCTACATTTATA**TAA**ctcgag**TGA**TACTACTAC | (*1*) |
| 22 | RT-PUT1Bot | GACAGTGACATTTTGTTTATTTGGGGTGTCATGGAAATATCTAAATTGAGTAGTAGTA**TCA**ctcgag**TTA**TATAAATGTAGC | (*1*) |
| 23 | RT-PUT2Top | CAGACATACATATTCATTTATAATGTTAAGATCAACTACTCGTAATACATTA**TAAATGA**ctcgagTACTAG | (*1*) |
| 24 | RT-PUT2Bot | GTGTGACGAATGATACTTGATGAAACTTTAGTATATCTAGTActcgag**TCATTTA**TAATGTATTAC | (*1*) |
| 25 | RT-PUT3Top | CATTCCTTCATTTACTTATATATAATCCGATTCTTGTACAATGGATTCACAAGAGCCT**TAA**ctcgag**TGA**AGAAAATTGC | (*1*) |
| 26 | RT-PUT3Bot | CAAGTGGAATGGTATCTGAATTAATTAATGCATTTGCAATTTTCT**TCA**ctcgag**TTA**AGGCTCTTG | (*1*) |
| 27 | 55-CgPUT1 | TTCTTTCCTGCGTTATCCCCTGATTCTGTGGATAACCGTACCATGGGATCGCCTGCAGAGATGTTAG | This work |
| 28 | 53-CgPUT1 | GAGGGGGGGCCCGGTACCCAATTCGCCCTATAGTGAGTCGCTTGTGATACTTGTGACGCTTG | This work |
| 29 | 35-CgPUT1 | TAGTGAGGGTTAATTGCGCGCTTGGCGTAATCATGGTCATCTGATGTCAAGACTCTTTACGCA | This work |
| 30 | 33-CgPUT1 | AACGCAGAAAATGAACCGGGGATGCGACGTGCAAGATTACCATA | This work |

| | | | |
|---|---|---|---|
| | | ACTTCATCTATCACACGCTTGTG | |
| 31 | 5C-CgPUT1 | CGAATAGTCCTCGAGAAACTGC | This work |
| 32 | 3C-CgPUT1 | GACCTTGTCTTTGGCAGTGAAG | This work |
| 33 | LOG-PUT1_fo | CTGATCGACAACTGCTCTAGAATC | This work |
| 34 | LOG-PUT1_re | GACGTTGTCTGACATACCTAGC | This work |
| 35 | 55-CgPUT2 | TTCTTTCCTGCGTTATCCCCTGATTCTGTGGATAACCGTACCATGG GCAGCACTCGAAACTCATAAC | This work |
| 36 | 53-CgPUT2 | GAGGGGGGGCCCGGTACCCAATTCGCCCTATAGTGAGTCGCCAAG GAATAGATCAGAAACAGAC | This work |
| 37 | 35-CgPUT2 | TAGTGAGGGTTAATTGCGCGCTTGGCGTAATCATGGTCATGATTT CAACTATCTACGCAGTGG | This work |
| 38 | 33-CgPUT2 | AACGCAGAAAATGAACCGGGGATGCGACGTGCAAGATTACCATC TTACACGACTGAGTGAACATGG | This work |
| 39 | 5C-CgPUT2 | GAAGCTACTACGGACACAACC | This work |
| 40 | 3C-CgPUT2 | GACATTTGGGAAGCGGTATGC | This work |
| 41 | LOG-PUT2_fo | CAGCTCTGATGGGTAACACTG | This work |
| 42 | LOG-PUT2_re | AGCTTGTTGAAACTTTGCTCG | This work |
| 43 | hk2 | CGTCAAGACTGTCAAGGAGGG | This work |
| 44 | hk3 | CATCATCTGCCCAGATGCGAAG | This work |
| 45 | YEp_ic_fwd | GTAATCTTGCACGTCGCATCC | This work |
| 46 | YEp_ic_rev | TACGGTTATCCACAGAATCAGGG | This work |
| 47 | SATflipp_fwd | CGACTCACTATAGGGCGAATTGG | This work |
| 48 | SATflipp_rev | TGACCATGATTACGCCAAGC | This work |
